# Going Against the Grain: Investigating the C4 wheat hypothesis with spatial transcriptomics

**DOI:** 10.64898/2025.12.11.691720

**Authors:** Tori Millsteed, David Kainer, Robert Sullivan, Xiaohuan Sun, Ka Leung Li, Likai Mao, Arlie Macdonald, Robert J Henry

**Affiliations:** Queensland Alliance for Agriculture and Food Innovation (QAAFI), University of Queensland, St Lucia 4072 QLD, Australia; ARC Centre of Excellence for Plant Success in Nature and Agriculture, University of Queensland, St Lucia 4072 QLD, Australia; Queensland Brain Institute, University of Queensland, St Lucia 4072 QLD, Australia; MGI Australia, Herston QLD 4006, Australia

**Keywords:** STOmics, seed, embryo, amino acids, carbon-nitrogen metabolism, grain filling, embryogenesis, dormancy

## Abstract

The possibility of a C4 photosynthetic pathway present in the developing grain of wheat, a C3 plant, has been the source of scientific debate. Wheat is critical to food security and may benefit greatly from the biological advantages conferred by C4 photosynthesis under heat and drought stress. Therefore, significant research has gone towards engineering wheat to use C4 biochemistry, resulting in the discovery of a unique photosynthetic pathway in the grain that has been suggested to be C4 specific. Here, we employed a spatial transcriptomics analysis of the developing wheat grain to further investigate the spatial expression patterns of C4 specific genes. Our results showed that most of the genes related to C4 photosynthesis were expressed in the grain in the theorised tissue locations, including phosphoenolpyruvate carboxylase (*ppc*) and pyruvate orthophosphate dikinase (ppdk). The photosynthetic pericarp cells were the site of *ppc* synthesis while the endosperm was the site of *ppc* carboxylation activity. Notably, isoforms of aspartate aminotransferase, alanine aminotransferase and malate dehydrogenase exhibited spatially distinct expression patterns, with tissue specificity, possibly linked to the unique functions of individual isoforms. As wheat performance under stress has been associated with the levels of expression of these C4 genes, confirmation of an active C4 pathway in the grain would have significant agronomic implications. Our results provide novel gene expression data for key genes related to photosynthesis, which could contribute to future development of highly productive, climate change resilient wheat varieties.

## Introduction

Wheat (*Triticum aestivum)* is one of the most consumed foods in the world and will continue to play an important role to ensure food security under climate change (Acevedo, 2018). Therefore, significant research has gone towards increasing wheat yields and stress tolerance. One avenue to do this is by enhancing photosynthetic efficiency, both in the leaves and non-leaf organs (Sanchez-Bragado *et al*, 2020; Aurus *et al*, 2021). Wheat is a C3 plant, meaning it utilises the C3 photosynthetic pathway, with ribulose-1,5-bisphosphate carboxylase/oxygenase (Rubisco) as the main CO_2_ fixing enzyme in the first step of photosynthesis. However, at higher temperatures, Rubisco loses efficiency for CO_2_ fixation, and increasingly fixes O_2_ instead, resulting in energetically wasteful photorespiration (Brautigam & Gowik, 2016). Therefore, heat stress represents a major limitation for productivity in C3 crops. This issue has been overcome by the evolutionarily younger C4 photosynthetic pathway observed in crops such as maize (*Zea mays*), sorghum (*Sorghum bicolor*) and sugar cane (*Saccharum officinarum*). C4 photosynthesis first emerged around 32-25 million years ago and has evolved independently many times (Sage *et al*, 2011). This pathway notably uses phosphoenolpyruvate carboxylase (*ppc*) instead of Rubisco in the first CO_2_ fixation step, eliminating the possibility of photorespiration (Williams *et al*, 2012), though Rubisco is still active in later steps. The vast majority of C4 plants also separate the light and dark reactions of photosynthesis into the mesophyll and bundle sheath cells, respectively, forming a structure known as Kranz anatomy. This allows for gas exchange in the mesophyll cells where *ppc* is active, and for carbon to be concentrated around Rubisco in the bundle-sheath cells for subsequent carbon fixation steps, preventing contact with O_2_ and maintaining Rubisco’s efficiency (Garner *et al*, 2016). However, C4 photosynthesis has also been reported to take place in some species without Kranz anatomy (Voznesenskaya *et al*, 2001; Sage, 2002). C4 photosynthesis confers significant biological advantages under heat and drought stress, therefore, the engineering of C3 crops such as wheat to utilise C4 biochemistry would be a major advance for agricultural production and food security (Cui, 2021).

Research for crop improvement has thus led to the major discovery of a possible C4 photosynthetic pathway active in the developing wheat grain. As early as 1976, C4-like metabolism was identified in the pericarp of barley (*Hordeum vulgare*) grains, based on the generation of the C4 photosynthetic product malic acid (Nutbeam & Duffus, 1976). This was followed by supporting findings in wheat that *ppc* activity was significantly higher in the ear compared to that of the leaves (Wirth *et al*, 1977; Singal *et al*, 1986), and the same was observed in rice (*Oryza sativa*) spikelets compared to the leaves (Imaizumi, *et al* 1990). Wheat ear gas exchange was also found to align with that of C3-C4 intermediate species (Ziegler-Jons *et al*, 1989), and ear photosynthesis was established to have greater tolerance to drought and heat stress than the leaves (Xu *et al*, 1990; Xu *et al*, 2003). However, these findings were then disputed by research in the 1990s and early 2000s. A radiolabelling study of ^14^CO_2_ suggested that developing wheat grains assimilated CO_2_ by the C3 pathway based on feeding the grains an external CO_2_ supply, with little fixation into the C4 acids malate and aspartate detected (Bort *et al*, 1995). Pericarp *ppc* was found to reassimilate CO_2_ already respired during grain filling, contributing to the greater stress tolerance of the ear (Bort *et al*,1996). Additionally, a study of water relations in the wheat ear showed that it responded metabolically to water stress in line with C3 plants (Tambussi *et al*, 2005). The same study assessed immunolocalization of Rubisco in the glumes, lemmas and awns, finding a lack of spatial segregation between mesophyll and bundle sheath cells and confirming the presence of C3 photosynthesis (Tambussi *et al*, 2005). However, the question of photosynthesis in the grain alone was not addressed.

The C4 wheat debate was then revived in 2016 by a transcriptomics analysis presenting new evidence for C4 photosynthesis in the developing wheat grain (Rangan *et al*, 2016). This study found that all genes required for an active C4 photosynthetic pathway were expressed in the grain at 14 days post anthesis (DPA), during the peak of grain filling. They were *ppc*, aspartate aminotransferase (*aat*), alanine aminotransferase (*gpt*), malate dehydrogenase (*mdh*), NAD-dependent malic enzyme (*me2*) and pyruvate orthophosphate dikinase (*ppdk*) (Rangan *et al*, 2016). In wheat and other grasses, these enzymes have multiple isoforms encoded by a number of isogenes which can differ in expression and activity. Therefore, the authors identified grain-expressed isoforms of each enzyme that could be C4-type and involved in photosynthesis in the grain. This was largely based on their subcellular localisation which differed from the localisation of the other, equivalent isoforms, and aligned with the C4 pathway (Rangan *et al*, 2016). Additionally, one grain-expressed isoform of *ppc* has an amino acid substitution associated with C4-type *ppc*s in known C4 grass species, and this was proposed to be a C4-type *ppc* unique to wheat. This isoform was also expressed at significantly higher levels in the grain than in the leaves, supporting the suggested role in grain photosynthesis (Rangan *et al*, 2016). Furthermore, this study pointed to chloroplast dimorphism between the tube- and cross-cell layers of the pericarp, similar to the structural differences between chloroplasts in mesophyll and bundle sheath cells in C4 plants. Thus, the developing wheat grain would be able to achieve spatial segregation of the photosynthetic reactions, reminiscent of Kranz anatomy (Rangan *et al*, 2016). And lastly, the study highlighted that wheat performance under stress has been associated with the levels of expression of these C4 genes (Jia *et al*, 2015; Zhang *et al*, 2019), emphasising the importance of characterising grain photosynthesis for future crop improvement efforts (Rangan *et al*, 2016).

However, their study was criticised for basing the model of C4 photosynthesis on transcriptomics data from the whole grain, lacking tissue specific detail, and overlooking previous evidence against C4 photosynthesis in the ear (Busch & Farquhar 2016; Hibberd & Furbank, 2016). Debate has continued about the merit of earlier studies as evidence for and against the different classifications of photosynthesis, and their implications on more recent findings (Henry *et al*, 2017; Henry *et al* 2020; Tambussi *et al* 2021). Addressing this, the authors published a follow-up study of tissue specific single-cell transcriptomics data (Pearce *et al*, 2015), analysing the differences between the outer pericarp, the inner pericarp and the endosperm (Rangan *et al*, 2024). Finding that *ppc* and *aat* were most highly expressed in the endosperm, the theorised model of C4 photosynthesis was revised to incorporate the endosperm tissue as the initial source of CO_2_ fixation (Rangan *et al*, 2024). That study represents the most detailed and up-to-date view of C4 photosynthesis in wheat, but limitations remain about the accuracy of single-cell transcriptomics (scRNA-seq) and the details that are lost during single-cell dissection, particularly in plant systems (Chen *et al*, 2023). In recent years scRNA-seq has been superseded by the development of spatial transcriptomics techniques, allowing for *in situ* capture and RNA-sequencing at subcellular resolution (Yin *et al*, 2023). This new approach to transcriptome analyses provides a more detailed view of gene expression across an entire tissue section and has recently been applied to a range of plant species and organs, including the developing wheat grain (Millsteed *et al*, 2025). Therefore, spatial transcriptomics could facilitate an updated analysis of C4 photosynthetic genes expressed in the grain and contribute new evidence to the C4 wheat debate. Here, we employed STOmics Stereo-seq to investigate the spatial expression patterns of genes related to C4 photosynthesis in the developing wheat grain, and their isoforms.

## Results

The STOmics spatial transcriptomics platform was used to measure spatial gene expression in the 14 DPA wheat seed, as described in Millsteed *et al* (2025). Seed sections were cut in a cryostat and mounted onto the STOmics assessment chips, allowing for *in situ* RNA capture and sequencing with spatial tags. The sequencing information was mapped to the IWGSC Chinese Spring RefSeq v2.1 reference genome (Zhu *et al*., 2021) and reconstituted back into the original spatial locations to generate a spatial gene expression matrix. The gene expression matrix was then overlaid with fluorescent microscope images of the original tissue, to place gene expression within the cell and tissue architecture. From this, key tissue and cellular groups could be identified and linked to regions of gene expression including the pericarp, endosperm, aleurone, crease and embryo (Figure 1). Due to challenges adhering the tissue sections to the chip surface, the most intact section was chosen as the focus of this study, though some folding of the tissue was present. Data from replicate chips have been used to support the findings where possible.

**Figure 1.**
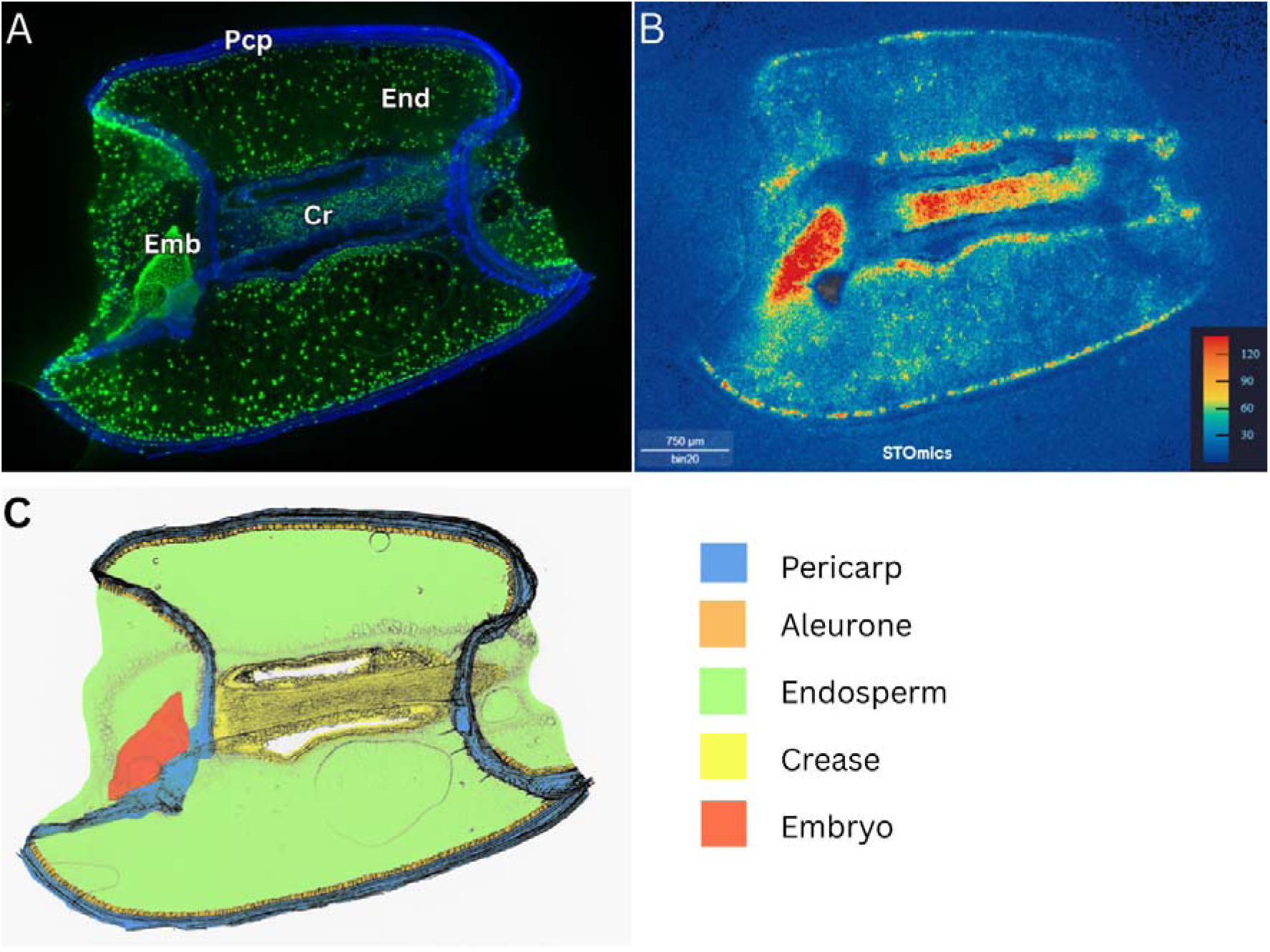
Fluorescent microscope image, gene expression matrix and diagrammatical representation of 14 days post anthesis (DPA) wheat grain. **A)** Fluorescent microscope image of wheat seed section, with cell walls in blue and cell nuclei in green, showing pericarp (Pcp), endosperm (End), embryo (Emb) and crease (Cr) tissues. **B)** Gene expression heat map of wheat seed section showing areas of highest gene expression in red and lowest gene expression in blue (in counts of transcripts). Scale bar = 750 μm. **C)** Diagrammatical representation of wheat seed section showing the pericarp, aleurone, endosperm, crease and embryo tissues represented by different colours, indicated in key.

Genes encoding the enzymes involved in C4 photosynthesis, and their isoforms, were identified and visualised in the gene expression matrix. Subgenome A, B and D homeologs were grouped together to capture the full expression of each isoform. Where genes were already named in the literature, those names have been used in this study. Where genes were unnamed, we assigned them identifiers based on chromosomal location, subcellular localisation (where chromosomal locations were the same) and decimal numbering (where chromosomal locations and subcellular localisation were the same) (Table S1).

### Phosphoenolpyruvate carboxylase (*ppc*)

The hexaploid wheat genome contains six isoforms of *ppc*, arwhich have been characterised for their expression in different organs of the wheat plant (Table S1) (Yamamoto *et al*, 2022). They are *ppc1a* (chr6), *ppc1b* (chr7), *ppc2* (chr5), *ppc3* (chr3), *ppc4* (chr3) and *ppc-b* (chr3). Three of these isoforms, *ppc1a*, *ppc2* and *ppc3,* are reported to be expressed in the developing grain (Yamamoto *et al*, 2014a; Yamamoto *et al*, 2022; Ruiz-Ballesta *et al*, 2016). In our analysis, all grain-expressed *ppc*s exhibited ubiquitous expression throughout the grain, lacking spatial specificity to a particular tissue or cell type (Figure S1). The isoforms reportedly not expressed in the grain were also investigated and their lack of expression confirmed (Table S1; Figure S1). Notably, *ppc3* has been suggested to be C4-type due to an amino acid substitution from an Arginine (R) residue to Serine (S) residue at position 884 in the protein sequence (Rangan *et al*, 2016). The R residue at this position is widely conserved in C3 *ppc*s, including all other wheat *ppc* isoforms, while substitutions such as Glycine (G) or S are observed in C4 *ppc*s (Figure 2) (Rangan *et al*, 2016). In the seed sections analysed, *ppc3* was expressed in both the endosperm and photosynthetic pericarp (Figure 2), which aligns with the proposed multi-cell model of C4 photosynthesis in the wheat grain (Table 1) (Rangan *et al*, 2024).

**Figure 2.**
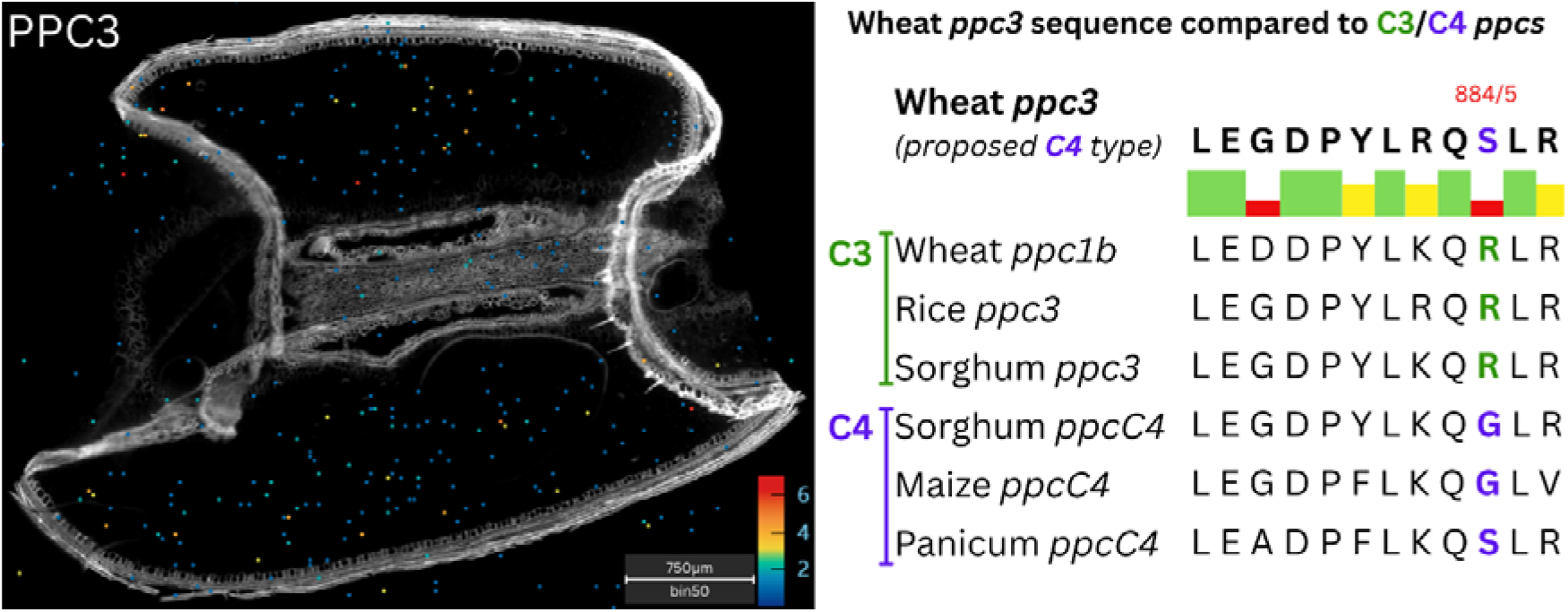
Proposed C4-type phosphoenolpyruvate carboxylase (*ppc3*) expression pattern and sequence information compared *ppc* sequences from other C3 and C4 species. Left) Gene expression matrix showing *ppc3* expression, where bins with highest gene expression are shown in red and those with lowest gene expression are shown in blue as per the heat map keys corresponding to each image (in transcript counts per bin). Scale bar = 750μm. **Right)** Amino acid sequence alignments of *ppc3* compared to other C3 *ppc*s (Wheat *ppc1b*, rice *ppc3* and sorghum *ppc3*) and known C4 *ppc*s (sorghum *ppcC4*, maize *ppcC4* and panicum *ppcC4*). Areas of sequence similarity are indicated by the green, yellow and red bars, while the amino acids at position 884/885 in the sequence are highlighted to show the differences between C3 and C4 *ppc*s.

**Table 1.** Comparison of expression values of genes suggested to be C4-specific in the developing wheat grain, from scRNA-seq data (Rangan *et al*, 2024) and spatial transcriptomics data (this study). Showing gene name, chromosome location (by chromosome number), RPKM expression values for the outer pericarp, inner pericarp and endosperm, reported in Rangan *et al*, 2024, and the spatial expression values in transcript counts per 1000 bins for the pericarp, endosperm, embryo and crease, reported in the present study.

| Gene Name | C4 genes chr. location | RPKM expression values (Rangan *et al*, 2024) |  |  | Spatial transcriptomics expression values (transcript counts per 1000 bins) |  |  |  |
| --- | --- | --- | --- | --- | --- | --- | --- | --- |
|  |  | Outer pericarp | Inner pericarp | End. | Pericarp | End. | Embryo | Crease |
| *ppc* | 3ABD (cyt) | 11 | 27 | 58 | 13 | 18 | 9 | 7 |
| *aat* | 3ABD (mt) | 539 | 1197 | 2134 | 30 | 109 | 243 | 175 |
|  | 7ABD (mt) | 32 | 91 | 63 | 9 | 6 | 16 | 3 |
| *gpt* | 2ABD (cyt) | 23 | 23 | 18 | 8 | 4 | 9 | 11 |
| *mdh* | 5ABD (mt) | 170 | 94 | 59 | 12 | 9 | 18 | 14 |
| *me2* | 2ABD (mt) | 51 | 26 | 7 | 6 | 5 | 16 | 11 |
| *ppdk* | 1ABD(chl) | 182 | 2404 | 11 | 119 | 7 | 18 | 2 |

### Aspartate aminotransferase (*aat*)

The spatial expression of four *aat* isoforms were investigated; *aat3* (chr3), *aat6(mt)* (chr6), *aat6(chl)* (chr6) and *aat7* (chr7). *aat3* was expressed ubiquitously and exhibited an order of magnitude greater overall expression than the other isoforms investigated (Table S3). This isoform was also more highly expressed in the embryo than any other tissue type (all adjusted approximate permutation tests p < 0.01). *aat3* was the suggested C4-type and, being strongly expressed in the endosperm, matched the spatial expression patterns proposed in the C4 wheat model (Table 1; Figure 3) (Rangan *et al*, 2024). Additionally, the *aat6(chl)* and *aat6(mt)* isoforms were more highly expressed in the embryo compared to the endosperm, (adjusted approximate permutation tests p < 0.01). The remaining isoforms were expressed throughout the grain at lower levels without specificity to a particular region or cell type (Figure S2).

**Figure 3.**
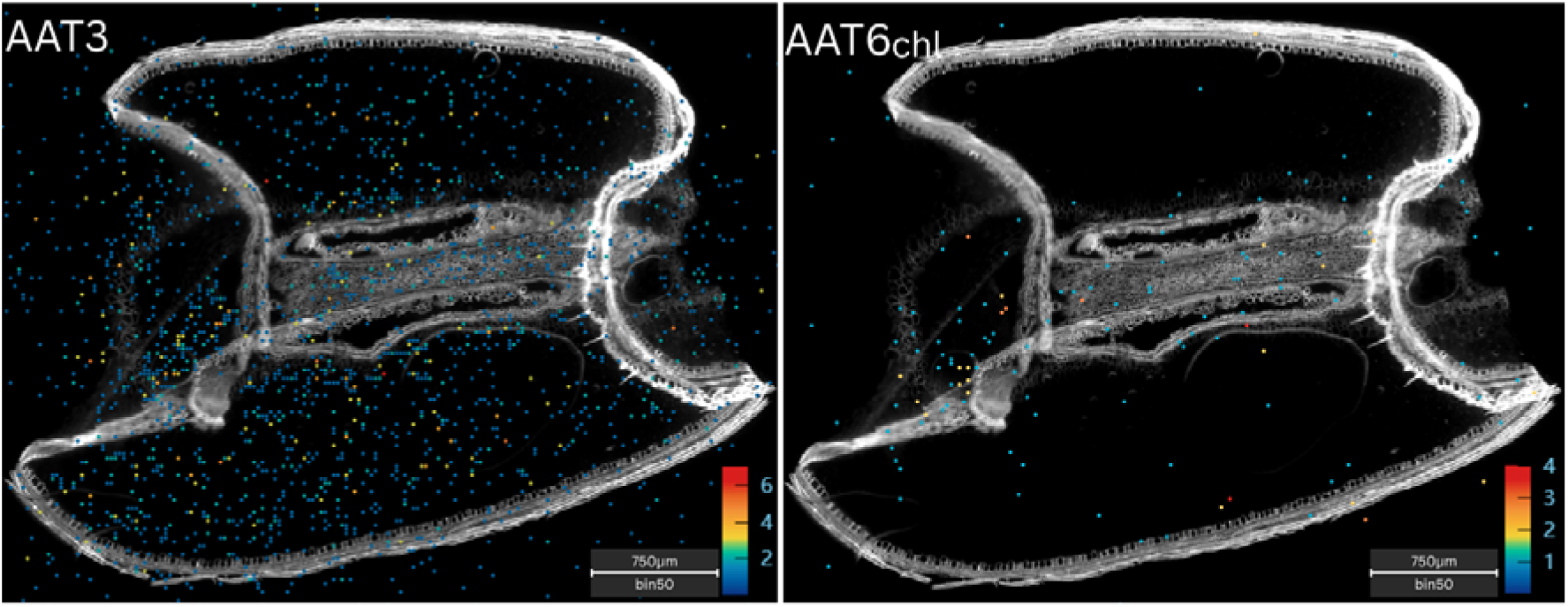
Gene expression matrices of two aspartate aminotransferase (*aat*) isoforms, *aat3* and *aat6(chl)*. The bins with highest gene expression are shown in red and those with lowest gene expression are shown in blue as per the heat map keys corresponding to each image (in transcript counts per bin). Scale bars = 750μm. **Left)** *aat3* gene expression pattern was more abundant than **Right)** *aat6(chl)* gene expression in all regions.

### Alanine aminotransferase (*gpt*)

There are four *gpt* isoforms characterised in the wheat genome: *gpt1* (chr1), *gpt2* (chr2), *gpt5* (chr5) and *gpt5Qsd1* (chr5). The *gpt1* isoform exhibited an order of magnitude greater overall expression than the other isoforms (Table S4) and was found to be more highly expressed in the endosperm compared to the crease (adjusted approximate permutation tests p < 0.01). The *gpt5Qsd1* isoform exhibited specificity to the embryo as it was more highly expressed in the embryo tissue than any other tissue type (all adjusted approximate permutation tests p < 0.01) (Figure 4). The *gpt2* isoform was the suggested C4-type and exhibited similar relative expression levels between the pericarp and endosperm, as was hypothesised in the C4 wheat model (Table 1) (Rangan *et al*, 2024).

**Figure 4.**
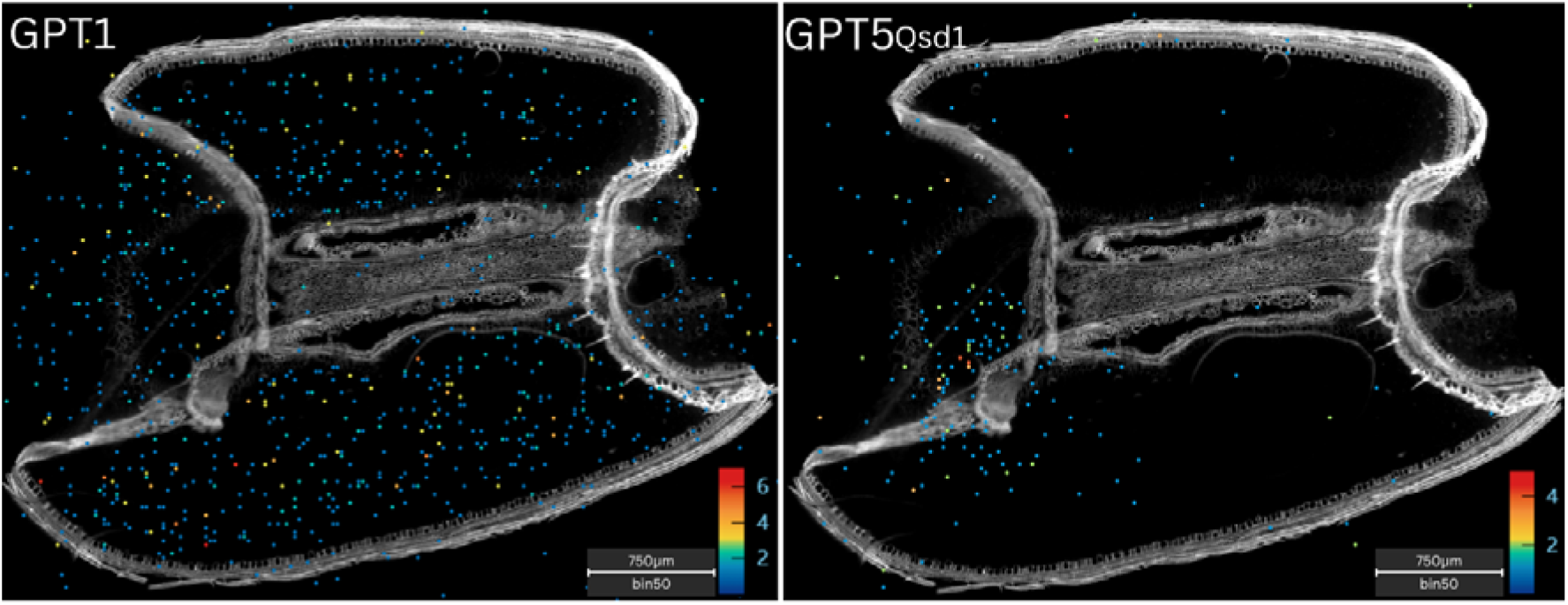
Gene expression matrices of two alanine aminotransferase (*gpt*) isoforms, *gpt1* and *gpt5Qsd1*. The bins with highest gene expression are shown in red and those with lowest gene expression are shown in blue as per the heat map keys corresponding to each image (in transcript counts per bin). Scale bars = 750μm. **Left)** *gpt1* was expressed throughout the endosperm and embryo while **Right)** *gpt5Qsd1* gene expression was specific to the embryo region.

### Malate dehydrogenase (*mdh*)

There are eight isoforms of *mdh* characterised in the wheat genome. They are *mdh1(mt)* (chr1), *mdh1(ct)* (chr1), *mdh1(chl)* (chr1), *mdh3* (chr3), *mdh5.1(gl)* (ch5), *mdh5.2(gl)* (chr5), *mdh7.1(chl)* (chr7), *mdh7.2(chl)* (chr7). An isoform which exhibited tissue specificity was *mdh1(mt)*, with significantly higher expression in the embryo compared to the other tissues (all adjusted approximate permutation tests p < 0.01) (Figure 5). The *mdh1(ct)* isoform was more highly expressed, especially in the endosperm, than the other isoforms, (Table 2) (Figure 5), while *mdh3* was found to be more highly expressed in the embryo, compared to the endosperm (adjusted approximate permutation tests p < 0.01) (Figure 5). *mdh5.1(gl)* was the isoform suggested to be C4-type and was expressed in all major tissues in the grain, exhibiting similar relative expression levels across all tissue types (Table 1).

**Figure 5.**
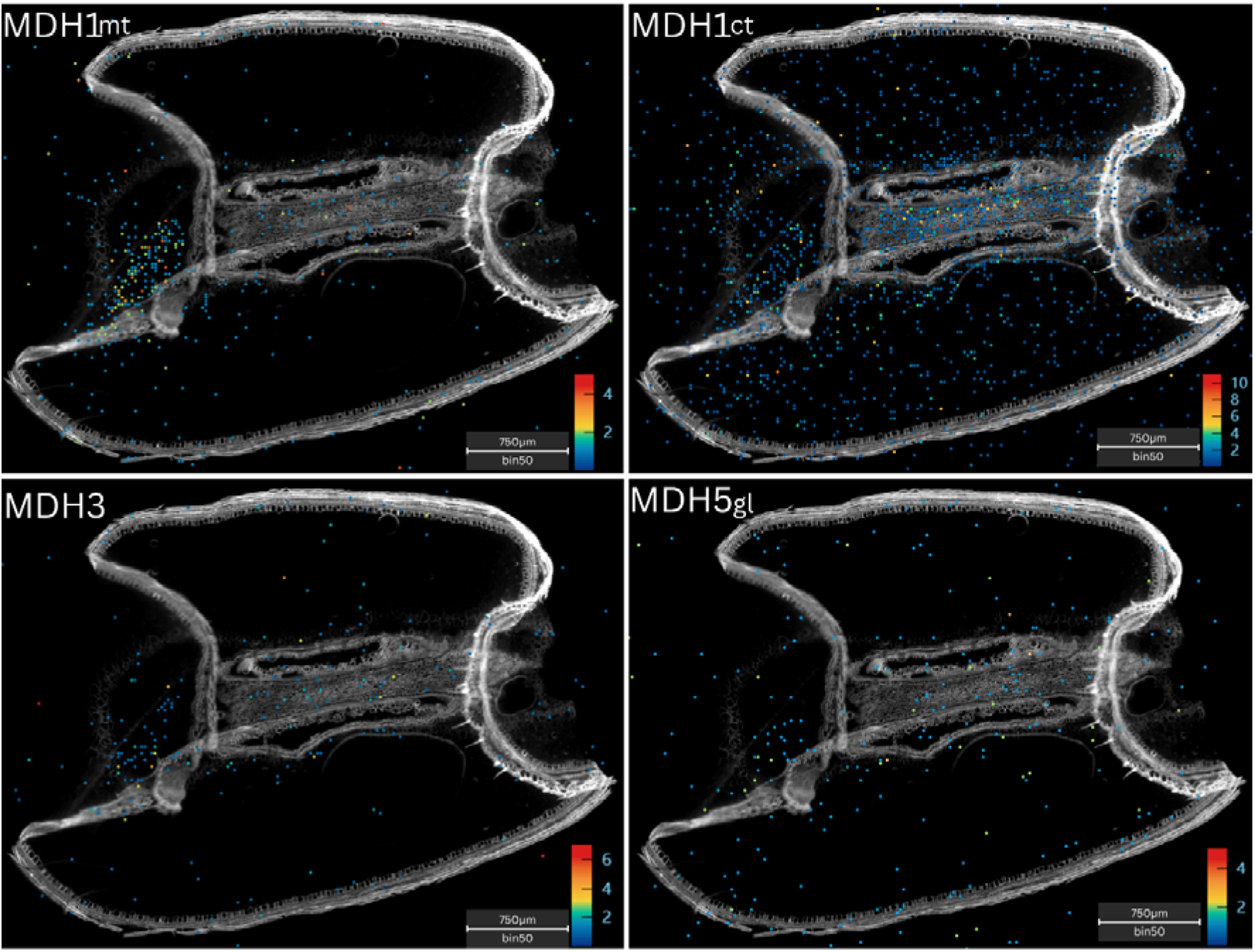
Gene expression matrices of four malate dehydrogenase (*mdh*) isoforms, *mdh1(mt), mdh1(ct), mdh3 and mdh5.1(gl).* The bins with highest gene expression are shown in red and those with lowest gene expression are shown in blue as per the heat map keys corresponding to each image (in transcript counts per bin). Scale bars = 750μm. **Top left)** *mdh1(mt)* gene expression pattern was highest in the embryo region compared to the **Top right)** *mdh1(ct)* gene expression pattern which as evident in all tissue types. **Lower left)** *mdh3* gene expression was also evident in the embryo region, whereas **Lower right)** *mdh5.1(gl)* gene expression was present in all tissues.

**Table 2.** Expression values of all C4-specific genes and their isoforms in transcript counts per 1000 bins, across the major tissue types in the developing wheat grain. Showing gene names of all grain expressed genes investigated, chromosome location (by chromosome number), and expression value in transcript count per 1000 bins for the pericarp, endosperm, embryo and crease.

| Gene name | Chromosome location | Spatial transcriptomics expression values in each tissue region (counts per 1000 bins) |  |  |  |
| --- | --- | --- | --- | --- | --- |
|  |  | Pericarp | Endosperm | Embryo | Crease |
| *ppc1a* | 6ABD | 17 | 5 | 22 | 13 |
| *ppc2* | 5ABD | 15 | 16 | 2 | 6 |
| *ppc3* | 3ABD | 13 | 18 | 9 | 7 |
| *aat3* | 3ABD | 30 | 109 | 243 | 175 |
| *aat(mt)* | 6ABD | 8 | 11 | 35 | 38 |
| *aat6(chl)* | 6ABD | 4 | 5 | 33 | 19 |
| *aatmt7* | 7ABD | 9 | 6 | 16 | 3 |
| *gpt1* | 1ABD | 6 | 53 | 32 | 16 |
| *gpt2* | 2ABD | 8 | 4 | 9 | 11 |
| *gpt5Qsd1* | 5ABD | 15 | 4 | 27 | 2 |
| *gpt5* | 5ABD | 3 | 3 | 12 | 2 |
| *mdh1(mt)* | 1ABD | 25 | 8 | 244 | 31 |
| *mdh1(ct)* | 1ABD | 30 | 91 | 261 | 78 |
| *mdh1(chl)* | 1ABD | 4 | 1 | 15 | 10 |
| *mdh3(mt)* | 3ABD | 14 | 8 | 77 | 18 |
| *mdh5.1(gl)* | 5ABD | 12 | 9 | 18 | 14 |
| *mdh5.2(gl)* | 5ABD | 10 | 3 | 4 | 11 |
| *mdh7.1(chl)* | 7ABD | 4 | 3 | 14 | 6 |
| *mdh7.2(chl)* | 7ABD | 5 | 2 | 11 | 5 |
| *me2(1)* | 1ABD | 2 | 2 | 1 | 4 |
| *me2(2)* | 2ABD | 6 | 4 | 16 | 11 |
| *ppdk1(ct)* | 1ABD | 88 | 28 | 112 | 13 |
| *ppdk1(chl)* | 1ABD | 119 | 7 | 18 | 2 |

### NAD-dependent malic enzyme (*me2*)

Two isoforms of the *me2* enzyme are found in wheat; *me2(1)* (chr1) and *me2(2)* (chr2). Both were expressed at relatively low levels in the seed sections analysed, therefore no significant difference in expression between tissues were identified. The *me2(2)* isoform was suggested to be the C4-type, playing a role in the photosynthetic pericarp tissue in the multi-cell model of C4 photosynthesis in the wheat grain (Rangan *et al*, 2024). However, this isoform didn’t show any specificity to the pericarp and even appeared to be expressed at lower levels in the pericarp compared to the endosperm, crease and embryo, though this couldn’t be confirmed statistically due to low overall expression (Table 1) (Figure 6).

**Figure 6.**
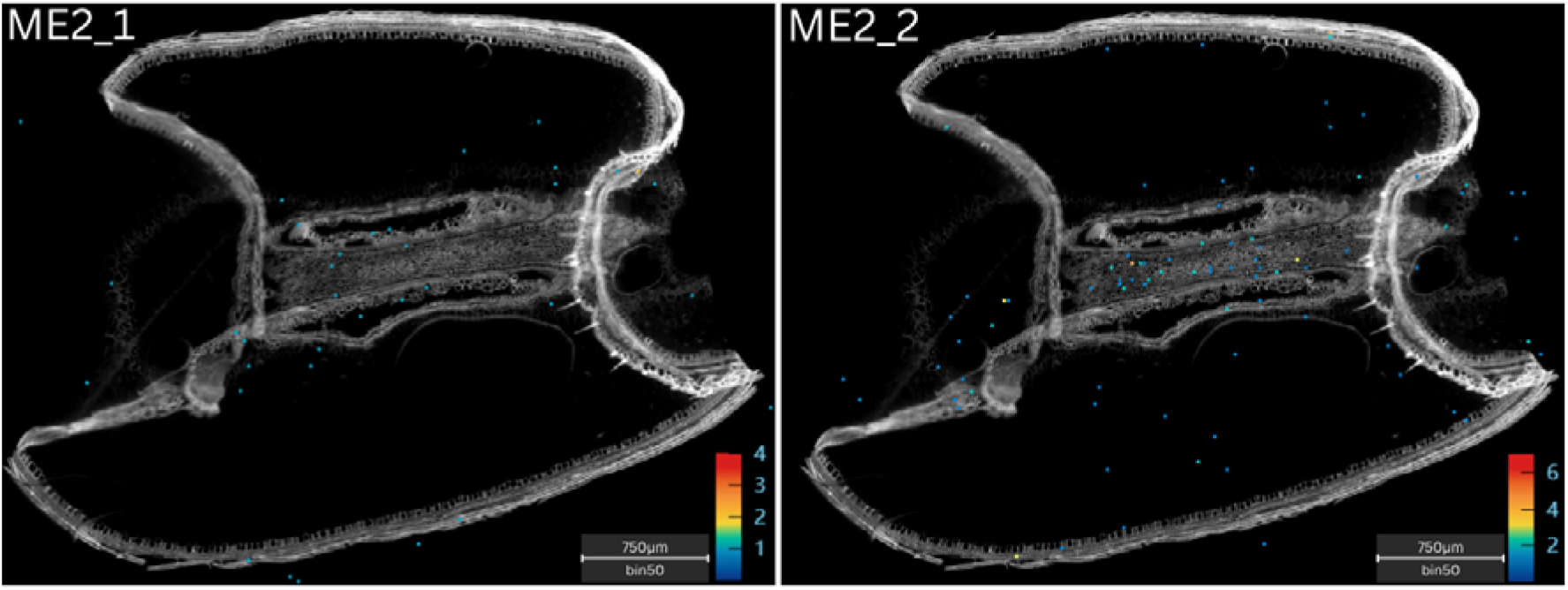
Gene expression matrices of two NAD-dependent malic enzyme (*me2*) isoforms, *me2(1)* and *me2(2)*. The bins with highest gene expression are shown in red and those with lowest gene expression are shown in blue as per the heat map keys corresponding to each image (in transcript counts per bin). Scale bars = 750μm. **Left)** *me2(1)* gene expression was low across all regions, while **Right)** *me2(2)* gene expression was evident in the embryo and crease.

### Pyruvate orthophosphate dikinase (*ppdk*)

Lastly, the two wheat *ppdk* isoforms, *ppdk1(ct)* (ch1) and *ppdk1(chl)* (ch1), were analysed. The *ppdk1(ct)* isoform showed strong tissue specificity with greater expression in the pericarp and embryo compared to the endosperm and crease (all adjusted approximate permutation tests p < 0.01) (Figure 7). This isoform also exhibited visibly greater overall expression levels compared to the *ppdk1(chl)* isoform. Though expressed at lower levels overall (Table S7), the *ppdk1(chl)* isoform showed the same pattern of tissue specificity to the pericarp (all adjusted approximate permutation tests p < 0.01) (Figure 7). Lastly, *ppdk1(chl)* was the suggested C4-type, and its expression aligned with the theorised pattern of being highly pericarp specific, supporting the multi-cell model of C4 photosynthesis in the wheat grain (Table 1) (Rangan *et al*, 2024).

**Figure 7.**
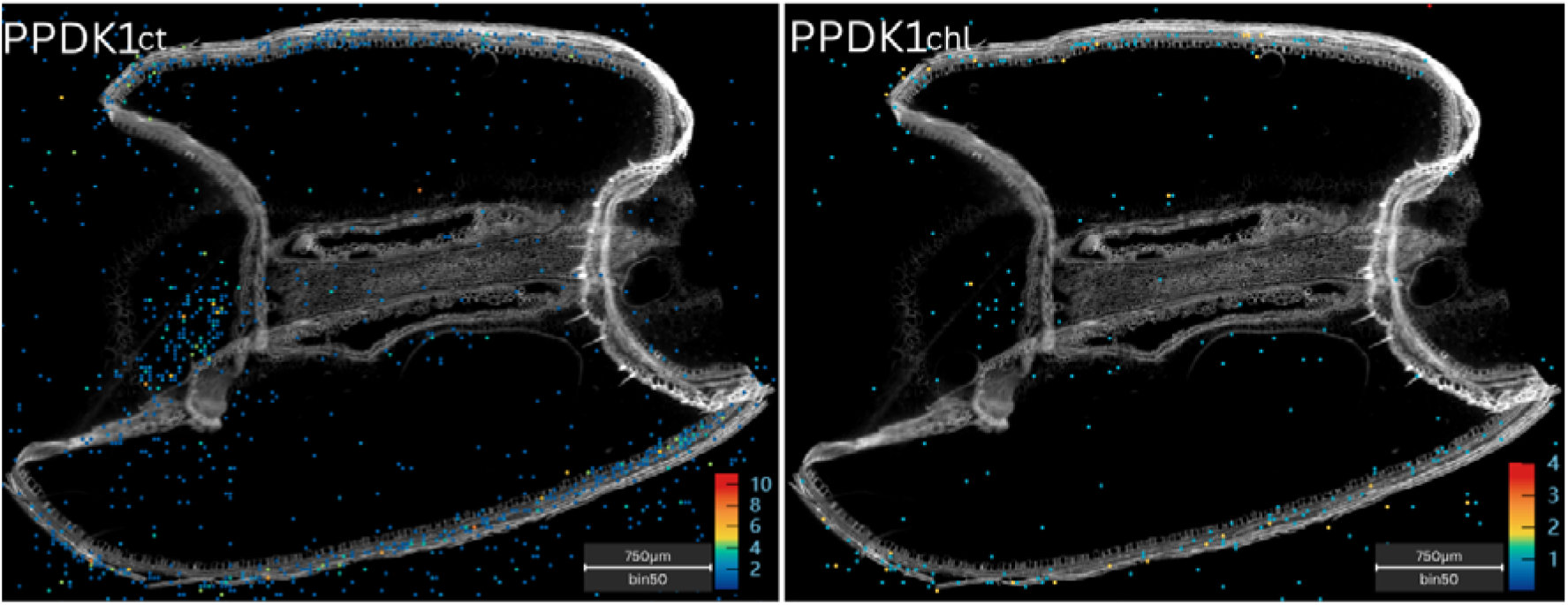
Gene expression matrices of two pyruvate orthophosphate dikinase (*ppdk*) isoforms, *ppdk1(chl)* and *ppdk1(ct).* The bins with highest gene expression are shown in red and those with lowest gene expression are shown in blue as per the heat map keys corresponding to each image (in transcript counts per bin). Scale bars = 750μm. The expression patterns of both **Left)** *ppdk1(ct)* and **Right)** *ppdk1(chl)* showed similar specificity to the pericarp and embryo, though *ppdk(ct)* was more highly expressed overall.

## Discussion

### Do these findings support the multi-cell model of C4 photosynthesis?

The suggested C4 isoforms of *ppc*, *aat*, *gpt* and *ppdk* all aligned with the spatial expression patterns hypothesised in the multi-cell model of C4 photosynthesis (Rangan *et al*, 2024). The C4-type *ppc* was expressed throughout the endosperm, where it may play a role in refixing already respired CO_2_ from respiration in the developing endosperm (Rangan *et al*, 2024). Due to an amino acid substitution from an R residue to a S residue, which isn’t observed in any other C3-type *ppc*, it is possible this isoform has a unique function related to grain photosynthesis. However, it should be noted that this isoform was also expressed at comparable levels in all tissue types and did not exhibit spatial specificity to the endosperm alone. Similarly, the suggested C4 isoform of *aat* was expressed in the endosperm and pericarp tissues, where it may play a role in the next step of a C4 photosynthetic pathway, shuttling carbon fixation products from the endosperm into the pericarp (Rangan *et al*, 2024). However, while highly expressed in the endosperm and pericarp, this *aat* isoform was found to be more highly expressed in the embryo tissue, raising questions about its role there. The C4-type *gpt* isoform was expressed in the pericarp as expected, where it could function to shuttle carbon and nitrogen products between the inner and outer pericarp for different steps in the C4 pathway. This *gpt* isoform was also expressed at comparable levels across all tissues in the seed. Lastly, *ppdk* was expressed in the expected tissue location, with significantly greater expression in the pericarp compared to any other tissue. The *ppdk* genes featured a strong pattern of expression in the inner pericarp region in the gene expression matrix, though our clustering method could not distinguish statistically between the inner and outer pericarp. In the pericarp tissue *ppdk* could be involved in the regeneration of *ppc* for the continuation of the C4 pathway (Rangan *et al*, 2024).

For the suggested C4 isoforms of *mdh* and *me2*, it was difficult to determine from the spatial data alone if their expression aligned with that proposed in the multi-cell model of C4 photosynthesis (Rangan *et al*, 2024). In NAD-type C4 photosynthesis, these genes work together to generate and then decarboxylate malate, creating a high concentration of CO_2_ around Rubisco in the bundle sheath cells (Ludwig, 2016). If *mdh* and *me2* undertake a similar role in the developing wheat grain, they would be expected to be specifically expressed in the photosynthetic pericarp tissue where Rubisco is active. However, expression of both the suggested C4 isoforms of *mdh* and *me2* appeared to be lower in the pericarp than other tissues within the grain, and no tissue specificity was identified for either. As some expression of these genes was observed in the pericarp, there remains the possibility that they take on a role in photosynthesis there, however, further targeted analysis is needed to confirm their function. While spatial transcriptomics represents a significant advancement in genetic research and offers new details about gene expression in the full tissue context, there remain limitations regarding genes expressed at very low levels or with high rates of transcript turnover. There are also limitations to verifying gene function based on expression location alone. Therefore, the spatial expression information must be considered in the context of multi-omics approaches and the wider knowledge base regarding target gene function.

### Roles of phosphoenolpyruvate carboxylase in C3 and C4 plants

The functional role of *ppc* is essential to the C4 wheat hypothesis because it represents the first major step in the C4 biochemical pathway. The spatial expression patterns of *ppc* isoforms identified in our data were in agreement with the suggested pathway in the wheat grain (Rangan *et al*, 2024), and earlier transcriptomics analyses (Pearce *et al*, 2015). However, there is growing evidence in the literature that *ppc3* takes on a role separate from photosynthesis. *ppc* is encoded by a small, multigene family that are conserved in C3 and C4 Poaceae species, including major crops such as wheat, rice (*Oryza sativa*), maize and sorghum (Yamamoto *et al*, 2022). These grasses usually contain 5-7 *ppc* isoforms, which have been characterised for their expression in specific plant organs. For example, *ppc1a* is expressed ubiquitously throughout the plant (Yamamoto *et al*, 2014a) while *ppc1b* is expressed specifically in the leaves (Yamamoto *et al*, 2022) and *ppc3* is specific to the seeds (Ruiz-Ballesta *et al*, 2016) and roots (Arias-Baldrich *et al*, 2017). The non-photosynthetic *ppcs* are thought to be involved in house-keeping functions, including supplying carbon skeletons to replenish the TCA cycle, carbon-nitrogen (C/N) metabolism, and pH homeostasis (Marin-Pena *et al*, 2024). In C4 species such as sorghum and maize, the seventh *ppc* isoform is *ppcC4* which catalyses the first carbon fixation step in C4 photosynthesis. In the Poaceae family, *ppcC4* likely evolved from *ppc1b* as a result of a chromosomal duplication event that took place approximately 45-60 million years ago (Yamamoto *et al*, 2022). As the ancestral form of the C4 *ppc* isoform, *ppc1b* has been found to be upregulated under osmotic stress conditions and as part of the plant nitrogen response. This isoform is retained in both C4 and most C3 Poaceae species but has been lost from the rice genome (Yamamoto *et al*, 2022).

Conversely, the *ppc* isoform suggested to be C4 specific in the developing wheat seed is *ppc3* (Rangan *et al*, 2016). This is the most abundant isoform in both the seeds and roots (Ruiz-Ballesta *et al*, 2016; Arias-Baldrich *et al*, 2017), and the only wheat *ppc* which features an S residue in place of an R residue at the amino acid position were R is widely conserved in C3 (non-photosynthetic) *ppcs* (Rangan *et al*, 2016). This substitution may be related to photosynthetic function and suggest a role in a C4-like photosynthetic pathway in the developing wheat seed, however, this does not explain the abundance of this isoform in the roots where there is no photosynthetic tissue. Grain expressed *ppc3* has long been thought to be involved in recapture of already respired CO_2_, supplying carbon to the TCA cycle in support of seed filling and development (Huppe and Turpin, 1994; Golombek et al, 1999). However, the complex roles of *ppc3* have recently been investigated in further detail in Arabidopsis (*Arabidopsis thaliana*) (Feria *et al*, 2022), wheat (Yamamoto *et al*, 2014c), rice (Yamamoto *et al*, 2015) and sorghum (Marin-Pena *et al*, 2024; de la Osa *et al*, 2025).

In Arabidopsis, mutant lines with knockouts of each individual *ppc* and *ppc* kinase isoform, were assessed for the impact on seed development and quality (Feria *et al*, 2022). All *ppc* mutants exhibited moderately reduced seed parameters, showing that the entire gene family was needed for normal growth and development (Feria *et al*, 2016). Additionally, the *ppc3* mutant showed significantly reduced global nitrogen, with a 39% reduction in protein content compared to the wildtype (Feria *et al*, 2022). Grain expressed *ppc* has also been linked to nitrogen metabolism in rice (Yamamoto *et al*, 2014b) and wheat (Yamamoto *et al*, 2014c), and the *ppc3* isoform was found to be co-expressed with fatty acid biosynthesis pathway genes in the developing rice seed. In rice, *ppc3* was also found to be localised in the aleurone and embryo of the developing seed (Yamamoto *et al*, 2015), differing from the spatial expression of *ppc3* in the wheat seed. In sorghum, knocking out the *ppc3* gene resulted in decreased germination rates under salinity stress, lower seed protein content and lower C/N ratios in the seed compared to the wildtype (de la Osa *et al*, 2025). *ppc3* silenced plants also showed greater sensitivity to ammonium nutrition at the roots, evidenced by the accumulation of NH4+ in the roots and deregulation of the normal TCA function (Marin-Pena *et al*, 2024). Therefore, *ppc3* appears to play an important role in nitrogen metabolism, pH regulation in both the seeds and roots, going beyond the classical view that the non-C4 *ppcs* primarily recycle carbon for the TCA cycle.

While our findings confirmed the expected spatial expression patterns of the *ppc3* isoform suggested by the multi-cell model of C4 photosynthesis (Rangan *et al*, 2024), there remains contrasting evidence regarding its role in the developing wheat grain. The expression of *ppc3* in the wheat seed along with other C4 specific genes, and its unique ‘non-C3’ amino acid substitution, are significant findings pointing to a role in photosynthesis, and warrant further investigation as a target pathway for improving grain development and yield. However, there is growing evidence that this *ppc* is involved with nitrogen metabolism and protein biosynthesis, critical to both seed development and the function of the root system. It is possible that *ppc3* takes on multiple roles in different tissues or that its role may differ between C3 and C4 species, which could explain the differing hypotheses about its function in wheat, rice and sorghum. Therefore, further research is still needed to validate the specific role of *ppc3* in the developing wheat grain, but all evidence is clear that this gene is critical to seed development and nutritional quality.

### Roles of malate dehydrogenase and NAD-dependent malic enzyme

In wheat *mdh* is represented by a multigene family containing 8 isogenes. The photosynthetic function of *mdh* in C4 plants is converting oxaloacetate into malate, which serves as a transport form of fixed carbon in the bundle sheath cells and contributes to the carbon concentrating mechanism around Rubisco (Ludwig, 2016). In C3 species, the *mdh* gene family is involved with a number of important reactions, including catalysing the conversion between oxaloacetate and malate, along with the interconversion of NADH and NAD⁺, which are vital for electron transport (Gietl, 1992; Imran *et al*, 2016). *mdh* and malate are also involved in regulating leaf respiration (Tomaz *et al*, 2010), embryo formation (Beeler et al, 2014) and root development (van der Merwe *et al*, 2008), as well as enhancing phosphorus uptake, facilitating nitrogen fixation, and improving tolerance to aluminium stress (Schulze *et al*, 2002). The variety of functions is unsurprising given the large number of isoforms, and is reflected in the distinct spatial expression patterns of our data. While the results didn’t indicate any *mdh* isoforms with spatial specificity to the pericarp tissue, this is in contrast with earlier scRNA-seq data showing that the suggested C4 *mdh* isoform was expressed at higher levels in the pericarp than the endosperm tissue (Pearce *et al*, 2015). The lack of pericarp specific expression in our data could be due to low overall expression, and the findings of some *mdh* expression in the pericarp means that a role in photosynthesis in the developing grain cannot be discounted. Further targeted analysis is needed to validate the function of the suggested C4 *mdh* isoform in wheat.

Only two *me2* isoforms have been identified in the wheat genome, one of which could take on a C4 photosynthetic role in the wheat grain (Rangan *et al*, 2016). In C4 photosynthesis, *me2* works with *mdh* in the bundle sheath cells, decarboxylating malate to supply CO_2_ to Rubisco. In C3 plants and non-photosynthetic tissues, *me2* catalyses the oxidative decarboxylation of malate to produce pyruvate, facilitating the exchange of carbon intermediates to maintain normal function of the tricarboxylic (TCA) cycle (Sun *et al*, 2019). This reaction contributes to photorespiration and energy metabolism, rather than directly to CO_2_ fixation as in C4 plants. The role of non-photosynthetic *me2* is not yet well understood but has been linked to increased abiotic stress tolerance, possibly due to balancing the concentration of malic acid in cells and reducing the production of radical oxygen species (Sun *et al*, 2019). It has been reported that both respiratory and C4-type *me2* are localised in the mitochondria of bundle sheath cells in the leaves of C4 species (Hudig *et al*, 2021), suggesting that relying on subcellular localisation or spatial expression patterns may not be sufficient to differentiate the functions of the C4 and non-C4 type. In our spatial data both *me2* isoforms were expressed throughout the grain with no spatial distinction between them, and neither with spatial specificity to the pericarp. The lack of specific expression in the photosynthetic tissue of the grain may indicate that neither isoform is involved in a photosynthetic pathway, however, the low overall expression makes this difficult to determine. Further research is still needed to validate if *me2* is involved in a C4 photosynthetic pathway in the developing wheat grain.

### Spatial expression differences identified between isoforms of C4 genes

Distinct spatial expression patterns between isoforms of *aat*, *gpt*, *mdh* and *ppdk* were reported for the first time in the present study, offering new insights about their unique roles in the developing wheat grain. Multiple isoforms of *aat* exhibited spatial specificity to the embryo tissue, including *aat3*, which was the suggested C4-type (Rangan *et al*, 2016). *aat* catalyses the reversible transamination between aspartate and 2-oxoglutarate, forming oxaloacetate and glutamate, a reaction that links the TCA cycle with amino acid biosynthesis (Han *et al*, 2021). This enzyme is thought to play an important role in embryo and endosperm development by integrating carbon and nitrogen metabolism to support rapid cell division and storage compound synthesis (Han *et al*, 2021). During embryo development, *aat* likely maintains a steady supply of aspartate-derived amino acids needed for growth (Sano *et al*, 2022), which is supported by its high expression in the embryo tissue. The aspartate amino acid family (leucine, threonine, methionine, isoleucine) have also been linked to embryo development in Arabidopsis, pea (Kirma *et al*, 2012) and rice (Sano *et al*, 2022). In its role in C4 photosynthesis, *aat* transports carbon fixation products from the mesophyll to bundle sheath cells and is hypothesised to play a similar role between the endosperm and pericarp tissue in the developing wheat grain (Rangan *et al*, 2024). However, the potential role of a C4-type *aat* in the embryo of the wheat grain is not yet characterised. The spatial transcriptomics data offers new detail about the expression of these genes during seed development, highlighting that *aat* likely plays an important role in embryo development.

Unique spatial expression pattens were also identified between the *gpt* isoforms, appearing to align with the already characterised functions of seed expressed *gpt* genes. (Agrawal et al, 2024). One *gpt* isoform identified as having embryo specific expression was known seed dormancy gene *TaQsd1* (*gpt5Qsd1*). In barley, the *Qsd1* gene encodes a mitochondrial *gpt* that regulates seed dormancy by modulating embryo metabolism during seed maturation (Sato *et al*, 2016). By converting alanine and 2-oxoglutarate into pyruvate and glutamate, *Qsd1* influences C/N metabolism and the energy balance in developing seeds (Sato *et al*, 2016). This gene is homologous with the *TaQsd1* genes in wheat, therefore strongly suggesting that the embryo specific expression supports its role in controlling seed dormancy (Onishi *et al*, 2017). Conversely, the *gpt1* isoform exhibited endosperm specific expression, suggesting a role related to endosperm development. Aside from seed dormancy, *gpt* genes have been linked to hypoxia tolerance and nitrogen use efficiency in a number of species (Agrawal et al, 2024). Additionally, the OsAlaAT1 gene in rice encodes a *gpt* that contributes to starch accumulation in the endosperm by regulating C/N metabolism during grain filling (Yang *et al*, 2015). Therefore, it’s possible that the *gpt1* isoform plays a role in starch biosynthesis during endosperm development. These results highlight the importance of spatial expression data to delineate functional differences between isogenes.

The *mdh* gene family also exhibited distinct expression patterns between isoforms. In particular, the *mdh1(mt)* and *mdh3* isoforms showed embryo specific expression. This result appears to be supported by earlier Arabidopsis studies which identified at least one *mdh* isoform with embryo specific function. Among the Arabidopsis *mdh* gene family, the plastid localised NAD-dependent isoform (At3g47520) is critical to embryo development, with knockouts of this gene found to be embryo-lethal (Schreier *et al*, 2018). Beyond the plastidial isoform, the mitochondrial isoforms of *mdh* (mMDH1 and mMDH2) in Arabidopsis support mitochondrial TCA-cycle and respiratory activity. Double knockout of these genes results in a substantial proportion of seeds arresting at the torpedo-embryo stage, highlighting their importance to late embryogenesis or seed maturation (Sew *et al*, 2016). Therefore, it naturally follows that isoforms with different sub-cellular localisation or tissue specific expression may take on different roles. Further, *mdh* isoforms with specific expression in the embryo likely take on roles specific to embryo development, while isoforms with ubiquitous expression may have more generalised, though still important, functions. It’s clear that a number of *mdh* genes are essential for optimal seed development, and the additional spatial expression data is a significant advancement in the knowledge about their individual roles. These new insights help distinguish which genes may be useful to enhance specific traits of interest and will facilitate more targeted genetic research.

Lastly, both *ppdk* isoforms showed spatial expression patterns localised to the photosynthetic pericarp tissue. The cytosolic (*ppdk1(ct))* isoform also exhibited specificity to the embryo tissue, and notably higher expression levels than the suggested C4, chloroplastic (*ppdk1(chl))* isoform. Whether by a C3 or C4 pathway, the high expression of *ppdk* genes in the pericarp supports photosynthetic and respiratory function through the conversion of pyruvate to *ppc*, which can then enter metabolic pathways such as gluconeogenesis and amino acid biosynthesis, or feed *ppc* reactions that refix CO_2_ (Chastain *et al*, 2011). Additionally, *ppdk*’s reversible reaction helps regenerate ATP and recycle carbon intermediates, making a significant contribution to grain filling (Zhang et al, 2021). In developing and germinating Arabidopsis embryos, *ppdk* is thought to supply *ppc* to support glycolysis and gluconeogenesis (Andriotis *et al*, 2010; Eastmond *et al*, 2015). Therefore, the embryo specific expression pattern of *ppdk* in the developing wheat grain could suggest a similar role that has not yet been characterised. The differences in expression levels between the two isoforms may be attributed to the number of genes encoding each. While the cytosolic isoform is encoded by three homologous genes, one from each subgenome in wheat, as is the case for most wheat genes, the chloroplastic isoform is only encoded by one gene from the A subgenome. These results highlight the utility of spatial transcriptomics to uncover new details about gene expression patterns, particularly between similar genes such as isoforms of the same gene family or subgenome homeologs.

## Conclusion

Implementing C4 biochemistry into C3 crops has long been explored as a way to enhance agronomic productivity, and confirmation of an active C4 pathway already present in the wheat grain would significantly advance this goal. Here, we provide new insight into the spatial expression of genes related to C4 photosynthesis and their isoforms. The expression of C4 isoforms of *ppc*, *aat*, *gpt* and ppdk aligned with the spatial organisation hypothesised in the multi-cell model of C4 photosynthesis (Rangan *et al*, 2024), suggesting that elements of a C4-like pathway may operate between the endosperm and pericarp tissues. However, the lack of clear tissue specificity of C4-type *mdh* and *me2* does not provide clear evidence to support the C4 wheat hypothesis. Additionally, the spatial data revealed novel and tissue-specific expression patterns for multiple isoforms, notably embryo specificity for some *aat*, *gpt*, *mdh* and *ppdk* genes. These results offer the detail to distinguish between the localisation and function of individual isoforms, which could facilitate more targeted genetic research in future. While spatial transcriptomics represents a powerful tool to investigate gene expression at detailed resolution, functional validation will be essential to determine the contribution of these genes to wheat carbon capture and subsequent yield.

## Materials and Methods

### Sample collection and STOmics stereo-seq experiment

Plant growth conditions, sample collection and the STOmics stereo-seq experiment were conducted as described in Millsteed *et al* (2025). In brief, seeds from 14 DPA wheat plants were sampled and embedded in Optimum Temperature Compound (OCT) and quickly frozen. The seeds were then cut using a cryostat (CryoStar Nx70, Epredia) to sections of 20µm thick, which were mounted and fixed onto the STOmics assessment chips. After tissue fixation, the sections were stained for fluorescent imaging by the Qubit ssDNA reagent, and fluorescent images were taken of each section using the 10× objective lens in both the FITC channel and DAPI channel, to record the original tissue and cell architecture. The samples were then permeabilised in 0.1% PR enzyme in 0.01M HCl buffer for 12 minutes at 37°C, so that the RNA could be released from the tissue sections and captured on the chips’ surface. Reverse transcription was then performed overnight at 37°C, and the next day the tissue was removed from the chips’ surface and the cDNA collected. The cDNA was then used for library construction by the Stereo-seq Library Preparation kit. The resulting libraries were pooled and sequenced by whole genome sequencing using the T7 PE100 flow cell. During the protocol, challenges arose with the tissue mounting step, resulting in some folding and breaking of the tissue sections. We proceeded with the most intact tissue sections for the subsequent analyses.

### SAW Pipeline

The SAW pipeline was undertaken according to Millsteed *et al* (2025). In brief, the SAW software suite (https://github.com/BGIResearch/SAW), with default parameters, was used to map the raw sequencing reads to the wheat reference genome IWGSC CS RefSeq v2.1 (Zhu *et al*., 2021). The coordinate identity (CID) of each read was used to place it spatially in its original position on the chips, and this data was overlayed with the fluorescent microscope images of the tissue sections before permeabilization, so that reads could be mapped to their spatial location within the tissue. The SAW software suite was then used to generate a matrix of expression count of all genes expressed at each spot on the chip. To handle the sparseness of the gene expression within each spot, adjacent spots could be grouped into different bin levels, such as Bin200 for 200×200 spots, for analysis at varying resolutions. It was determined that Bin50 was the ideal resolution for this data. Once the gene expression matrices were generated, the chip containing the most intact replicate was designated as the focus for the subsequent analysis, while the remaining chips were referenced to provide replication where possible.

### Identification of gene expression clusters

Gene expression clusters were identified using the methods described in Millsteed *et al* (2025). In brief, the gene expression matrices were filtered to remove bins with low counts and background noise, while retaining bins within the tissue boundary. The expression data was normalised by the log1p method (Booeshaghi & Pachter, 2021) and then underwent principal component analysis (PCA). A neighbourhood graph of bins was then generated using the PCA representation of the expression matrix, followed by a spatial neighbours graph. These graphs were then embedded into two dimensions for ease of visualisation using UMAP.

Leiden clustering was then undertaken using the spatial neighbours data. This network analysis tool grouped bins based on the expression profile of the genes at each location. All transcripts captured on the chip were included in the network, resulting in 11-12 distinct gene expression clusters being identified for each chip. Overlaying the spatial Leiden clusters with the fluorescent microscope images of the tissue allowed us to link eight clusters to functional cellular groups, and these clusters were subsequently used to measure differential expression of specific genes between different regions of the grain.

### Investigation of photosynthetic genes

The spatial expression patterns of genes suggested to be involved in C4-type photosynthesis in the developing wheat grain were analysed, as well as the expression of the genes encoding the relevant isoenzymes. These included six *ppc* isoforms, four *aat* and *gpt* isoforms, eight *mdh* isoforms, and two *me2* and *ppdk* isoforms. To differentiate between isoforms, they were each uniquely named. Where genes were already named in the literature, those names have been used in this study. Where genes were unnamed, we assigned them identfiers based on chromosomal location, subcellular localisation (where chromosomal locations were the same) and decimal numbering (where chromosomal locations and subcellular localisation were the same).

The spatial expression of each isoform was visualised in the gene expression matrix using the StereoMap app. The homologous subgenome A, B and D genes encoding each isoenzyme were grouped together to capture the full expression of each. Furthermore, a sub-sample of 25% of bins from each cluster were randomly selected and used to undertake differential expression analyses. The bins were selected from clusters generated using the log1p normalised expression data, and those bin IDs were then used to extract the unnormalized MID count data from those locations on the chip.

The raw count data for the photosynthetic target genes at each randomly sampled bin ID (location) were then used for differential expression analysis to quantify the differences in expression of isoenzymes across different tissue types. The previously generated spatial gene expression clusters were used to indicate boundaries between different tissue regions.

To determine the statistical significance of the differences in gene expression we used an approximate (Monte Carlo) permutation test. This test was chosen as the highly zero-inflated spatial expression data would violate the normality assumption of a standard t-test and produce poor results under an equivalent rank-based test. In the approximate permutation test, 1,000,000 random permuted samples of the two sets of gene expression data being tested (ie bins from two different tissues) were generated, which provided an estimate of the p-value. Binomial theory was then applied to give a 99.9 % confidence interval for the true p-value. The estimated p-value confidence intervals were then adjusted by a Bonferroni multiple testing correction factor of 468, the total number of tests conducted. Resultingly, we have a high level of confidence that the true adjusted p-values were less than the upper limits of the adjusted confidence intervals, as reported in Tables S8-S10.

## Acknowledgements

This project was supported by the ARC Centre of Excellence for Plant Success in Nature and Agriculture (CE 200100015).

## Author contributions

RH conceived and supervised the project. TM, RS, KXS and KLL conducted the experiments. TM, DK, LM and AM conducted the analysis. TM wrote the manuscript with input from DK, RS, AM and RH. All authors read the final version of the paper.

## Data availability

The high-throughput data that supports the findings of this study are openly available in Gene Expression Omnibus at https://www.ncbi.nlm.nih.gov/geo/query/acc.cgi?acc=GSE298021, accession number GSE298021. The scripts used have been uploaded to GitHub (https://github.com/tori-millsteed/Wheat_seed_spatial_transcriptomics_Millsteed2025).

## Supporting information

Supporting data file 1: *supporting_data_1_additional_tables_and_figures.docx*

Supporting data file 2: *supporting_data_2_permutation_test_results.xlsx*

Supporting data file 3: *supporting_data_3_permutation_test_results_table_legends.docx*

