## Supporting data 1 for "Going Against the Grain: Investigating the C4 wheat hypothesis with spatial transcriptomics"

**Table S1.** **Phosphoenolpyruvate carboxylase sequence information.** Showing gene ID, gene name, chromosomal location, expression location and amino acid sequence from position 874 to 886, highlighting the amino acid substitution that is conserved across C3 *ppc*s, but differs for the grain expressed *ppc3* isoform.

| **Gene ID** | **Gene Name** | **Chromosome** | **Expression location** | **AA sequence 874-886** |
| --- | --- | --- | --- | --- |
| TraesCS6A03G0505900 | PPC1a_A | 6A | Roots/seeds | LEGDPYLKQ**R**LR |
| TraesCS6B03G0615400 | PPC1a_B | 6B | Roots/seeds | LEGDPYLKQ**R**LR |
| TraesCS6D03G0441600 | PPC1a_D | 6D | Roots/seeds | LEGDPYLKQ**R**LR |
| TraesCS7A03G0847000 | PPC1b_A | 7A | Leaves | LEDDPYLKQ**R**LR |
| TraesCS7B03G0663800 | PPC1b_B | 7B | Leaves | LEDDPYLKQ**R**LR |
| TraesCS7D03G0790600 | PPC1b_D | 7D | Leaves | LEDDPYLKQ**R**LR |
| TraesCS5A03G0481000 | PPC2_A | 5A | Ubiquitous | LEGDPYLKQ**R**LR |
| TraesCS5B03G0479900 | PPC2_B | 5B | Ubiquitous | LEGDPYLKQ**R**LR |
| TraesCS5D03G0444100 | PPC2_D | 5D | Ubiquitous | LEGDPYLKQ**R**LR |
| TraesCS3A03G0734400 | PPC3_A | 3A | Seeds | LEGDPYLRQ**S**LR |
| TraesCS3B03G0841200 | PPC3_B | 3B | Seeds | LEGDPYLRQ**S**LR |
| TraesCS3D03G0678400 | PPC3_D | 3D | Seeds | LEGDPYLRQ**S**LR |
| TraesCS3A03G0316700 | PPC4_A | 3A | Leaves | LESDPYLRQ**R**LL |
| TraesCS3B03G0401400 | PPC4_B | 3B | Leaves | LESDPYLRQ**R**LL |
| TraesCS3D03G0317900 | PPC4_D | 3D | Leaves | LESDPYLRQ**R**LL |
| TraesCS3B03G0014500 | PPCb_B | 3B | (Bacterial ppc) | SANNRSLRR**L**IE |
| TraesCS3D03G0011300 | PPCb_d | 3D | (Bacterial ppc) | SANNRSLRR**L**IE |

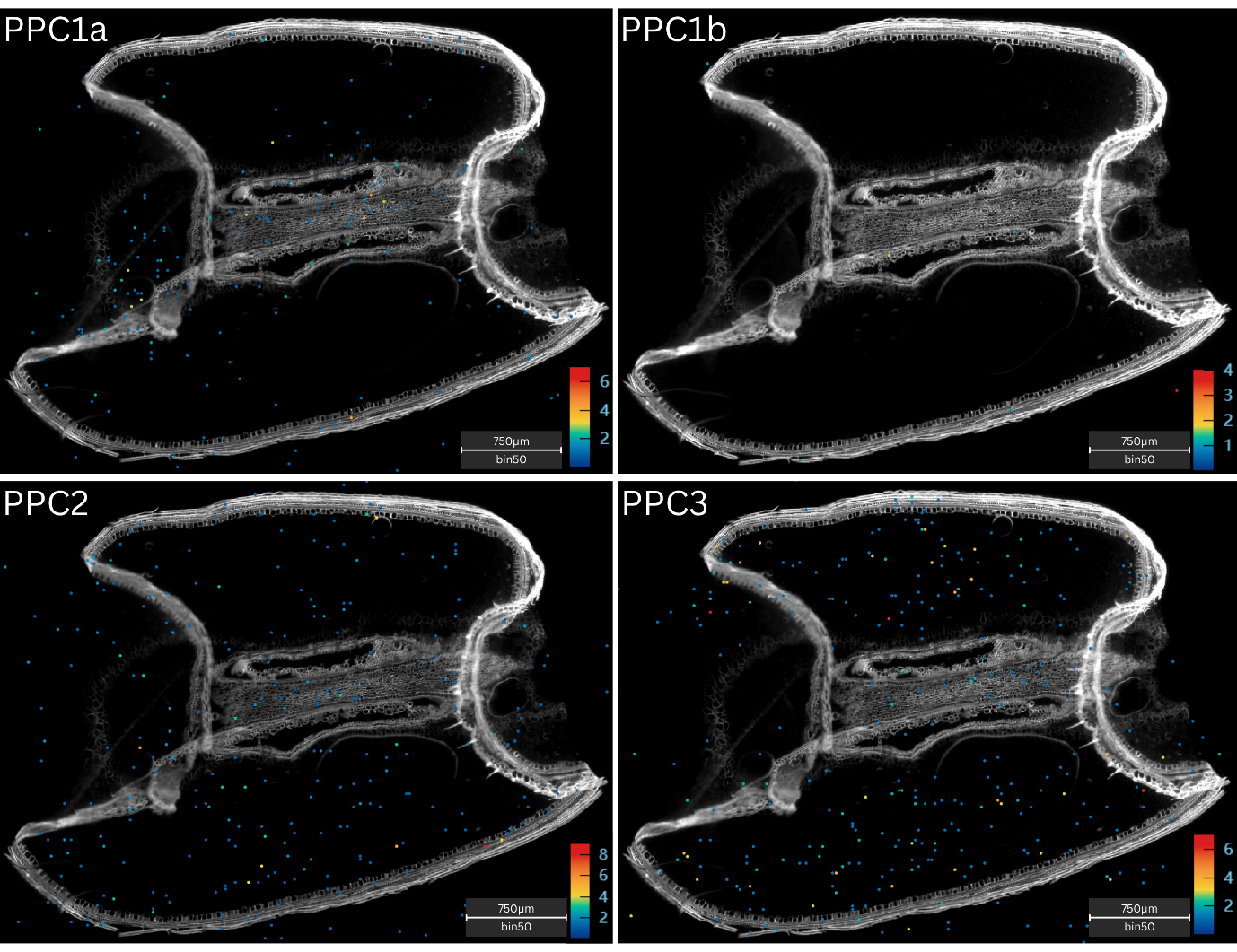

**Figure S1. Spatial gene expression patterns of phosphoenolpyruvate carboxylase isoforms in the 14 days post anthesis (DPA) wheat grain.** The bins with highest gene expression are shown in red and those with lowest gene expression are shown in blue as per the heat map keys corresponding to each image (in transcript counts per bin). Scale bars = 750μm.

**Table S2. Phosphoenolpyruvate carboxylase genes overall expression across replicate chips.** Showing gene ID, gene name, chromosomal location and overall expression in transcript counts for the replicate chips analysed.

| **Gene ID** | **Gene Name** | **Chromosome** | **Overall expression in transcript counts** | | |
| --- | --- | --- | --- | --- | --- |
|  |  |  | **Chip 1** | **Chip 2** | **Chip 3** |
| TraesCS6A03G0505900 | PPC1a_A | 6A | 154 | 209 | 130 |
| TraesCS6B03G0615400 | PPC1a_B | 6B | 104 | 105 | 50 |
| TraesCS6D03G0441600 | PPC1a_D | 6D | 161 | 205 | 123 |
| TraesCS7A03G0847000 | PPC1b_A | 7A | 10 | 17 | 6 |
| TraesCS7B03G0663800 | PPC1b_B | 7B | 12 | 6 | 7 |
| TraesCS7D03G0790600 | PPC1b_D | 7D | 2 | 7 | 3 |
| TraesCS5A03G0481000 | PPC2_A | 5A | 191 | 460 | 266 |
| TraesCS5B03G0479900 | PPC2_B | 5B | 346 | 812 | 461 |
| TraesCS5D03G0444100 | PPC2_D | 5D | 329 | 837 | 399 |
| TraesCS3A03G0734400 | PPC3_A | 3A | 231 | 302 | 147 |
| TraesCS3B03G0841200 | PPC3_B | 3B | 406 | 1013 | 719 |
| TraesCS3D03G0678400 | PPC3_D | 3D | 313 | 505 | 309 |
| TraesCS3A03G0316700 | PPC4_A | 3A | 0 | 0 | 0 |
| TraesCS3B03G0401400 | PPC4_B | 3B | 0 | 0 | 0 |
| TraesCS3D03G0317900 | PPC4_D | 3D | 0 | 0 | 0 |
| TraesCS3B03G0014500 | PPCb_B | 3B | 0 | 0 | 0 |
| TraesCS3D03G0011300 | PPCb_d | 3D | 0 | 0 | 0 |

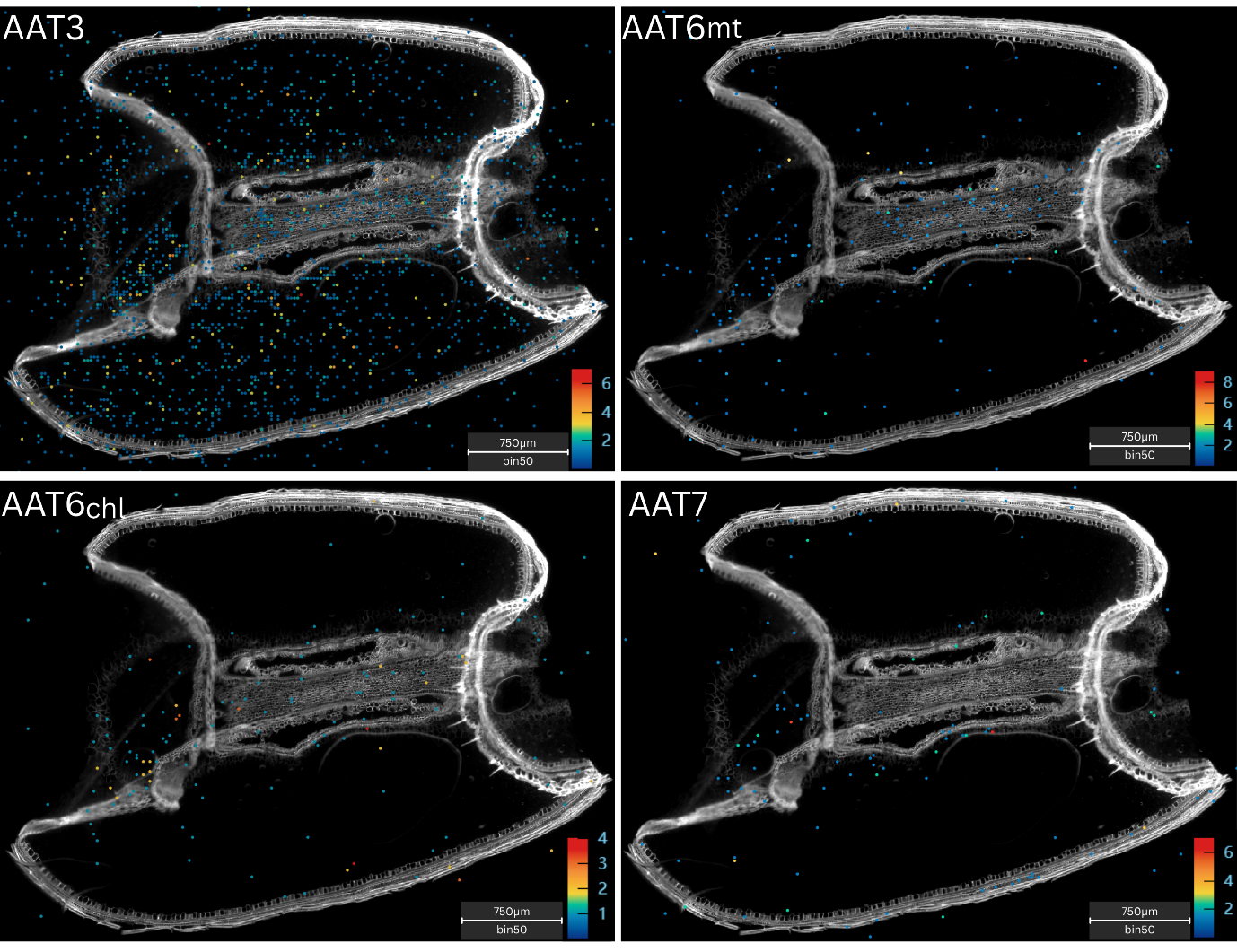

**Figure S2. Spatial gene expression patterns of aspartate aminotransferase isoforms in the 14 days post anthesis (DPA) wheat grain.** The bins with highest gene expression are shown in red and those with lowest gene expression are shown in blue as per the heat map keys corresponding to each image (in transcript counts per bin). Scale bars = 750μm.

**Table S3. Aspartate aminotransferase** **genes overall expression across replicate chips.** Showing gene ID, gene name, chromosomal location and overall expression in transcript counts for the replicate chips analysed.

| **Gene ID** | **Gene Name** | **Chromosome** | **Overall expression in transcript counts** | | |
| --- | --- | --- | --- | --- | --- |
|  |  |  | **Chip 1** | **Chip 2** | **Chip 3** |
| TraesCS3A03G0737300 | AAT_ct3A | 3A | 1913 | 6433 | 3507 |
| TraesCS3B03G0844300 | AAT_ct3B | 3B | 1211 | 3959 | 1908 |
| LOC123079235 | AAT_ct3D | 3D | 2212 | 6960 | 3853 |
| TraesCS7A03G0864200 | AAT_mt7A | 7A | 79 | 176 | 71 |
| TraesCS7B03G0751700 | AAT_mt7B | 7B | 97 | 106 | 30 |
| TraesCS7D03G0872300 | AAT_mt7D | 7D | 130 | 408 | 115 |
| TraesCS6A03G0924100 | AAT_chl6A | 6A | 149 | 104 | 76 |
| TraesCS6B03G1111000 | AAT_chl6B | 6B | 158 | 198 | 94 |
| TraesCS6D03G0797000 | AAT_chl6D | 6D | 138 | 155 | 78 |
| TraesCS6A03G0040500 | AAT_mt6A | 6A | 176 | 163 | 89 |
| TraesCS6B03G0060300 | AAT_mt6B | 6B | 134 | 127 | 110 |
| TraesCS6D03G0046000 | AAT_mt6D | 6D | 235 | 166 | 166 |

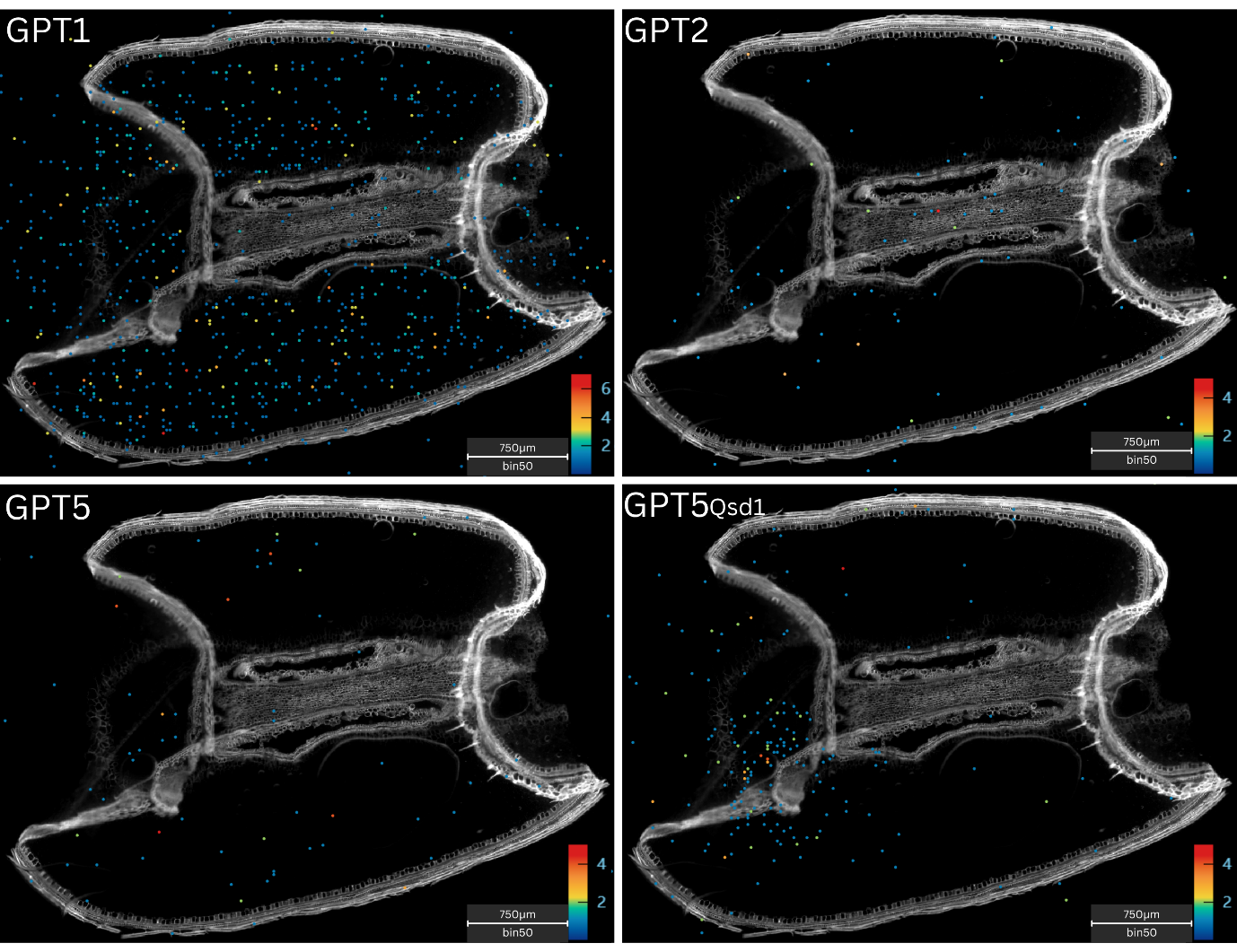

**Figure S3. Spatial gene expression patterns of alanine aminotransferase isoforms in the 14 days post anthesis (DPA) wheat grain.** The bins with highest gene expression are shown in red and those with lowest gene expression are shown in blue as per the heat map keys corresponding to each image (in transcript counts per bin). Scale bars = 750μm.

**Table S4. Alanine aminotransferase** **genes overall expression across replicate chips.** Showing gene ID, gene name, chromosomal location and overall expression in transcript counts for the replicate chips analysed.

| **Gene ID** | **Gene Name** | **Chromosome** | **Overall expression in transcript counts** | | |
| --- | --- | --- | --- | --- | --- |
|  |  |  | **Chip 1** | **Chip 2** | **Chip 3** |
| TraesCS2A03G0347900 | GPT_2A | 2A | 77 | 71 | 59 |
| TraesCS2B03G0467200 | GPT_2B | 2B | 72 | 92 | 65 |
| TraesCS2D03G0365600 | GPT_2D | 2D | 75 | 128 | 81 |
| TraesCS5A03G0558200 | GPT_5AQsd1 | 5A | 90 | 28 | 29 |
| TraesCS5B03G0568200 | GPT_5BQsd1 | 5B | 155 | 226 | 119 |
| TraesCS5D03G0523900 | GPT_5CQsd1 | 5D | 198 | 99 | 80 |
| TraesCS5A03G0151800 | GPT_5A | 5A | 97 | 264 | 143 |
| TraesCS5B03G0163300 | GPT_5B | 5B | 18 | 10 | 6 |
| TraesCS5D03G0171800 | GPT_5D | 5D | 71 | 210 | 75 |
| TraesCS1A03G0207600 | GPT_1A | 1A | 1402 | 4637 | 2032 |
| TraesCS1B03G0275700 | GPT_1B | 1B | 396 | 535 | 251 |
| TraesCS1D03G0200400 | GPT_1D | 1D | 1013 | 3016 | 1171 |

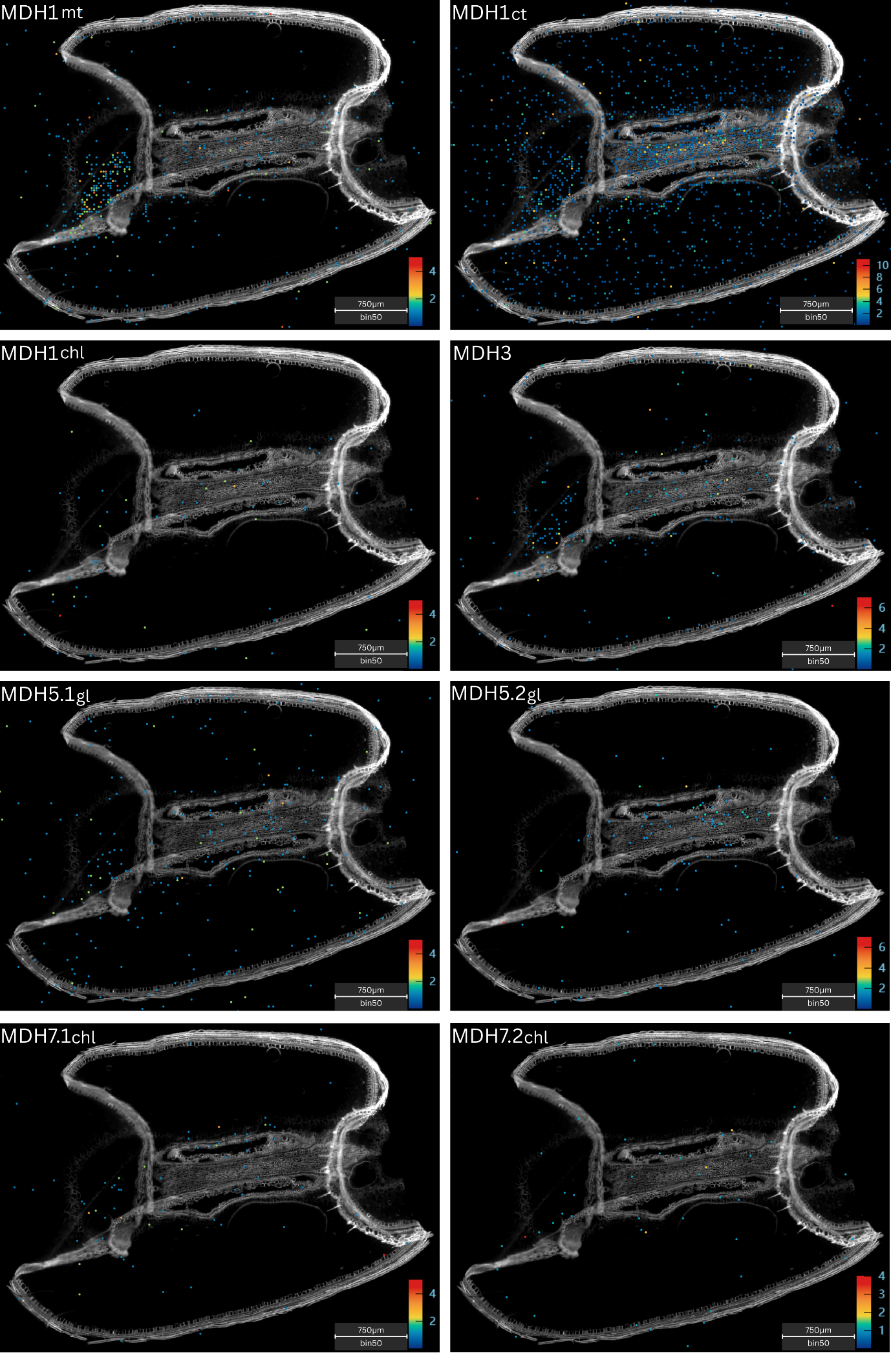

**Figure S4. Spatial gene expression patterns of malate dehydrogenase isoforms in the 14 days post anthesis (DPA) wheat grain.** The bins with highest gene expression are shown in red and those with lowest gene expression are shown in blue as per the heat map keys corresponding to each image (in transcript counts per bin). Scale bars = 750μm.

**Table S5. Malate dehydrogenase** **genes overall expression across replicate chips.** Showing gene ID, gene name, chromosomal location and overall expression in transcript counts for the replicate chips analysed.

| **Gene ID** | **Gene Name** | **Chromosome** | **Overall expression in transcript counts** | | |
| --- | --- | --- | --- | --- | --- |
|  |  |  | **Chip 1** | **Chip 2** | **Chip 3** |
| TraesCS1A03G1008400 | MDH_mt1A | 1A | 275 | 163 | 120 |
| TraesCS1B03G1190400 | MDH_mt1B | 1B | 499 | 609 | 254 |
| TraesCS1D03G0971000 | MDH_mt1D | 1D | 387 | 408 | 174 |
| TraesCS1A03G0416600 | MDH_ct1A | 1A | 2062 | 4554 | 1819 |
| TraesCS1B03G0509100 | MDH_ct1B | 1B | 1131 | 1513 | 585 |
| TraesCS1D03G0398800 | MDH_ct1D | 1D | 1786 | 3516 | 1585 |
| TraesCS1A03G0854400 | MDH_chl1A | 1A | 54 | 97 | 54 |
| TraesCS1B03G0988300 | MDH_chl1B | 1B | 57 | 106 | 56 |
| TraesCS1D03G0824900 | MDH_chl1D | 1D | 67 | 74 | 38 |
| TraesCS3A03G0605900 | MDH_mt3A | 3A | 114 | 43 | 14 |
| TraesCS3B03G0686400 | MDH_mt3B | 3B | 320 | 280 | 108 |
| TraesCS3D03G0557700 | MDH_mt3D | 3D | 220 | 283 | 88 |
| TraesCS5A03G0033000 | MDH_gl5A | 5A | 199 | 626 | 270 |
| TraesCS5B03G0026800 | MDH_gl5B | 5B | 226 | 713 | 342 |
| TraesCS5D03G0043100 | MDH_gl5D | 5D | 166 | 386 | 275 |
| TraesCS5A03G0967000 | MDH_cr5A | 5A | 83 | 75 | 35 |
| TraesCS5B03G1017200 | MDH_cr5B | 5B | 61 | 82 | 39 |
| TraesCS5D03G0920800 | MDH_cr5D | 5D | 56 | 48 | 31 |
| LOC123154897 | MDH_chl7A | 7A | 39 | 13 | 4 |
| TraesCS7B03G0546000 | MDH_chl7B | 7B | 67 | 37 | 14 |
| TraesCS7D03G0651600 | MDH_chl7D | 7D | 48 | 16 | 15 |
| TraesCS7A03G0934200 | MDH_ub7A | 7A | 27 | 46 | 20 |
| TraesCS7B03G0783200 | MDH_ub7B | 7B | 49 | 70 | 40 |
| TraesCS7D03G0901200 | MDH_ub7D | 7D | 35 | 71 | 27 |

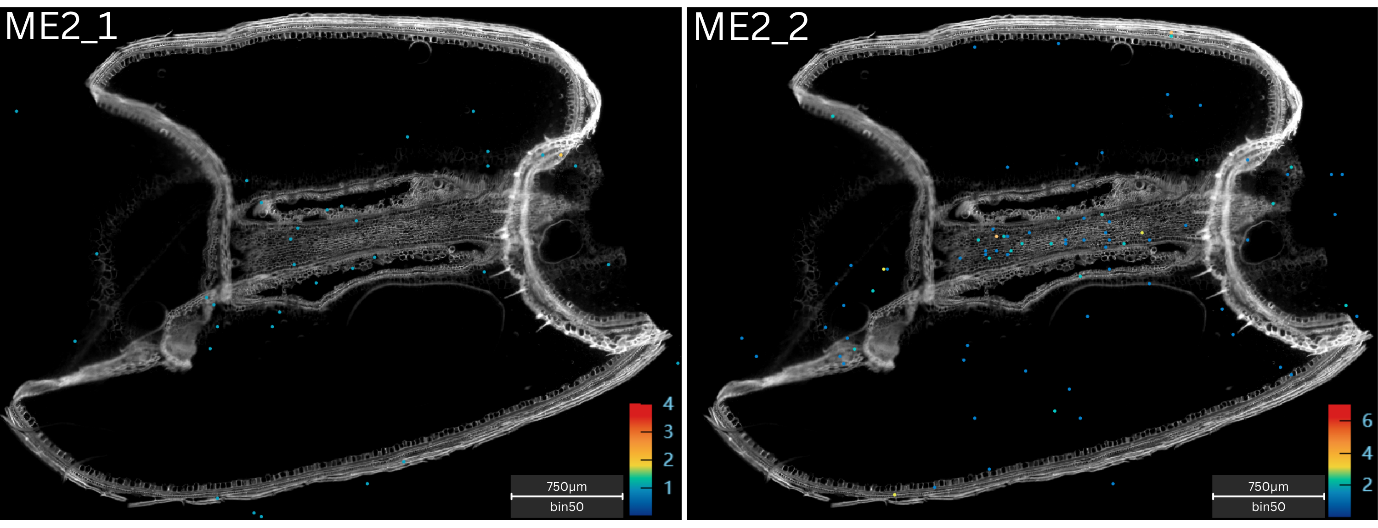

**Figure S5. Spatial gene expression patterns of NAD-dependent malic enzyme isoforms in the 14 days post anthesis (DPA) wheat grain.** The bins with highest gene expression are shown in red and those with lowest gene expression are shown in blue as per the heat map keys corresponding to each image (in transcript counts per bin). Scale bars = 750μm.

**Table S6. NAD-dependent malic enzyme** **genes overall expression across replicate chips.** Showing gene ID, gene name, chromosomal location and overall expression in transcript counts for the replicate chips analysed.

| **Gene ID** | **Gene Name** | **Chromosome** | **Overall expression in transcript counts** | | |
| --- | --- | --- | --- | --- | --- |
|  |  |  | **Chip 1** | **Chip 2** | **Chip 3** |
| TraesCS2A03G0540800 | ME2_2A | 2A | 84 | 65 | 53 |
| TraesCS2B03G0633800 | ME2_2B | 2B | 92 | 65 | 22 |
| TraesCS2D03G0499700 | ME2_2D | 2D | 71 | 47 | 24 |
| TraesCS1A03G0302700 | ME2_1A | 1A | 42 | 77 | 32 |
| TraesCS1B03G0387000 | ME2_1B | 1B | 13 | 5 | 3 |
| TraesCS1D03G0293000 | ME2_1D | 1D | 69 | 66 | 67 |

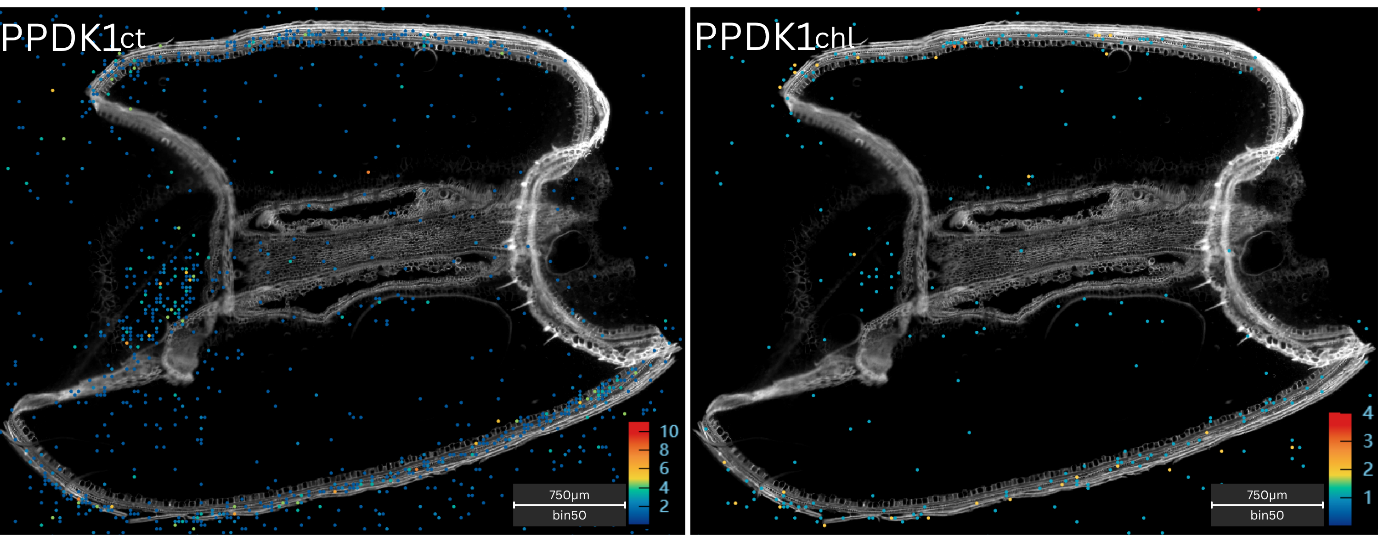

**Figure S6. Spatial gene expression patterns of pyruvate orthophosphate dikinase isoforms in the 14 days post anthesis (DPA) wheat grain.** The bins with highest gene expression are shown in red and those with lowest gene expression are shown in blue as per the heat map keys corresponding to each image (in transcript counts per bin). Scale bars = 750μm.

**Table S7. Pyruvate orthophosphate dikinase** **genes overall expression across replicate chips.** Showing gene ID, gene name, chromosomal location and overall expression in transcript counts for the replicate chips analysed.

| **Gene ID** | **Gene Name** | **Chromosome** | **Overall expression in transcript counts** | | |
| --- | --- | --- | --- | --- | --- |
|  |  |  | **Chip 1** | **Chip 2** | **Chip 3** |
| TraesCS1A03G0652200 | PPDK_chl1A | 1A | 697 | 2080 | 725 |
| TraesCS1A03G0652400 | PPDK_ct1A | 1A | 1380 | 3111 | 1038 |
| TraesCS1B03G0741000 | PPDK_ct1B | 1B | 609 | 1658 | 579 |
| TraesCS1D03G0610200 | PPDK_ct1D | 1D | 1418 | 2625 | 884 |
