## Supporting data 3 for "Going Against the Grain: Investigating the C4 wheat hypothesis with spatial transcriptomics"

Table legends for permutation test results (Tables S8-S10)

**Table S8. Chip 1 permutation test results:** Approximate permutation tests comparing the expression of grouped homologous gene triplicates across different tissue types (spatial gene expression clusters) to determine significant differences in spatial expression. This table represents data analysed from Chip 1 (ID: D02266B1), and includes the common names of the gene triplicates to be analysed, the names of the cluster groups to be compared, the observed difference in means, the estimated p-value, the adjusted p-value (by a Bonferroni multiple testing correction factor of 468), the number of replicates used by the permutation test, the gene names from the reference genome, ie the gene IDs, and the cluster IDs.

**Table S9. Chip 2 permutation test results:** Approximate permutation tests comparing the expression of grouped homologous gene triplicates across different tissue types (spatial gene expression clusters) to determine significant differences in spatial expression. This table represents data analysed from Chip 2 (ID: D02266A4), and includes the common names of the gene triplicates to be analysed, the names of the cluster groups to be compared, the observed difference in means, the estimated p-value, the adjusted p-value (by a Bonferroni multiple testing correction factor of 468), the number of replicates used by the permutation test, the gene names from the reference genome, ie the gene IDs, and the cluster IDs.

**Table S10. Chip 3 permutation test results:** Approximate permutation tests comparing the expression of grouped homologous gene triplicates across different tissue types (spatial gene expression clusters) to determine significant differences in spatial expression. This table represents data analysed from Chip 3 (ID: D02263D4), and includes the common names of the gene triplicates to be analysed, the names of the cluster groups to be compared, the observed difference in means, the estimated p-value, the adjusted p-value (by a Bonferroni multiple testing correction factor of 468), the number of replicates used by the permutation test, the gene names from the reference genome, ie the gene IDs, and the cluster IDs.
